# A pair of E3 ubiquitin ligases compete to regulate filopodial dynamics and axon guidance TRIM67 regulates filopodial stability and axon guidance

**DOI:** 10.1101/529222

**Authors:** Nicholas P. Boyer, Laura E. McCormick, Fabio L. Urbina, Stephanie L. Gupton

## Abstract

Appropriate axon guidance is necessary to form accurate neuronal connections. Guidance cues stimulate reorganization of the cytoskeleton within the distal growth cone at the tip of the extending axon. Filopodia at the periphery of the growth cone have long been considered sensors for axon guidance cues, yet how they perceive and respond to extracellular cues remains ill-defined. Our previous work found that the filopodial actin polymerase VASP is regulated via TRIM9-dependent nondegradative ubiquitination, and that appropriate VASP ubiquitination and deubiquitination are required for axon turning in response to the guidance cue netrin-1. Here we show that the TRIM9-related protein TRIM67 antagonizes VASP ubiquitination by outcompeting the TRIM9:VASP interaction. This antagonistic role is required for netrin-1 dependent filopodial responses, axon turning and branching, and fiber tract formation. We suggest a novel model that coordinated regulation of nondegradative VASP ubiquitination by a pair of ligases is a critical element of axon guidance.

## INTRODUCTION

Axon guidance toward appropriate synaptic partners is critical to the formation of the intricate neuronal networks found in mature organisms. Extracellular guidance cues direct axon navigation and are sensed by transmembrane receptors and interpreted through downstream signaling pathways (Kolodkin and Tessier-lavigne, 2011). Receptors often localize to the tips of actin rich filopodial protrusions in the axonal growth cone, which is a dynamic, cytoskeleton-rich structure at the distal end of extending axons (Shekarabi and Kennedy, 2002). Directionally biased remodeling and movement of growth cones, coupled with progressive elongation and condensation of axons, produces the turning behavior in axon guidance (Plachez and Richards, 2005). The localization of signals within the growth cone to allow for this directional bias requires tight regulation of effectors, such as cytoskeletal remodeling proteins (Dent et al., 2011). However, the mechanisms allowing for highly localized regulation in the growth cone are not understood, especially those that allow for rapid alteration of protein function and cytoskeletal dynamics in response to extracellular cues.

The guidance cue netrin-1 and its receptor DCC are required for midline-crossing behavior of many CNS axons (Bin et al., 2015; Fazeli et al., 1997; Serafini et al., 1996), including those in the corpus callosum of the placentalian brain (Fothergill et al., 2014). There is a large body of work demonstrating the function of signaling and cytoskeletal proteins activated during responses to netrin-1 (Boyer and Gupton, 2018). For example, the Ena/VASP actin polymerases are critical to filopodia formation and maintenance in neurons (Dent et al., 2007; Kwiatkowski et al., 2007), particularly downstream of DCC in netrin-1 signaling (Lebrand et al., 2004). Recent work has also shown that negative regulation of downstream effectors prior to netrin-1 signaling is required for appropriate netrin response (Menon et al., 2015; Plooster et al., 2017). For example, the E3 ubiquitin ligase tripartite motif protein 9 (TRIM9) is required for non-degradative ubiquitination and inhibition of vasodilator stimulated phosphoprotein (VASP) (Menon et al., 2015). VASP ubiquitination negatively impacts filopodia stability. This metric of filopodia stability is an element of growth cone response to extracellular cues like netrin-1 (Dent et al., 2007; Gupton and Gertler, 2007; Lebrand et al., 2004). Loss of VASP ubiquitination is necessary for growth cone filopodial response to netrin-1 (Menon et al., 2015), however the factors that reduce VASP ubiquitination or increase deubiquitination in the presence of netrin are unknown; furthermore regulators of TRIM9 have not been identified.

Tripartite motif protein 67 (TRIM67) is a class 1 TRIM protein along with TRIM9, sharing identical domain organization and 63.3% sequence identity (Short and Cox, 2006). Our recent work described a line of mice lacking *Trim67* and showed that TRIM67 is required *in vivo* for the appropriate development of several axons tracts including the netrin-sensitive corpus callosum (Boyer et al., 2018). Additionally, we found that TRIM67 interacts with both TRIM9 and the netrin-1 receptor DCC. Little is known about the cellular function of TRIM67, though a previous study reported TRIM67-dependent ubiquitination of 80K-H, a negative regulator of a Ras protein, in neuroblastoma cells (Yaguchi et al., 2012). However, no role has been described for TRIM67 in the regulation of axon guidance.

Here we describe a surprising, antagonistic role for TRIM67 in the ubiquitination of VASP in murine embryonic cortical neurons, and demonstrate that appropriate regulation of VASP ubiquitination is required for filopodial and axonal responses to netrin-1. We demonstrate that TRIM67 interacts with the actin polymerase VASP and negatively regulates its TRIM9-dependent ubiquitination. We provide evidence that this antagonism occurs via TRIM67 competitively inhibiting the interaction between TRIM9 and VASP. Using a combination of cell biological and biochemical approaches, we show that genetic deletion of *Trim67* results in increased VASP ubiquitination and basal defects in filopodia dynamics, as well as a loss of acute filopodial and growth cone responses to netrin. Additionally, netrin-dependent axon turning and branching responses are impaired by loss of *Trim67*. We extend these *in vitro* findings into the animal, where we find a perinatal delay in the developmental completion of the netrin-sensitive corpus callosum. These experiments suggest a model: a pair of closely related TRIM proteins generate a “yin and yang” like modulation of VASP function in axonal growth cones that allows filopodia to effectively search their environment, providing fidelity in netrin-1 dependent axon guidance.

## RESULTS

### TRIM67 is involved in netrin-dependent axon guidance

We recently generated mice carrying a *Trim67* allele flanked by loxP sites (*Trim67^Fl/Fl^*). Germline deletion of *Trim67* resulted in a thinner and smaller corpus callosum in the adult (Boyer et al., 2018). In light of this phenotype, we examined the perinatal development of the callosum in newborn (P0) and 4-day old (P4) *Trim67^+/+^*: Nex-Cre:Tau-Lox-STOP-Lox-GFP and Trim67^Fl/Fl^:Nex-Cre:Tau-Lox-STOP-Lox-GFP littermates. The Nex promoter drives Cre expression and recombination in excitatory, postmitotic cortical and hippocampal neurons starting at embryonic day (E) 11.5 (Goebbels et al., 2006). We analyzed corpus callosum development in serial sections (**Fig.1A**) at P0. Deletion of *Trim67* widened the interhemispheric distance between leading callosal fibers (**Fig.1A**, arrows) 80, 160, and 240 μm posterior to the last midline-crossing section of the corpus callosum (**Fig.1B**, p = 0.0055; p = 0.008; p = 0.045, respectively), suggesting there was a delay in axon extension toward the midline in *Trim67^Fl/Fl^* brains. Concordantly, corpus callosum growth was delayed toward the posterior of the brain at P0 (length of corpus callosum from fornix; *Trim67^+/+^* 453±22 μm, *Trim67^Fl/Fl^* 360±25 μm; p = 0.0342). By P4, when the callosum has completed midline crossing (Wahlsten, 1984), the callosum extended a shorter distance caudally in *Trim67^Fl/Fl^* littermates (**Fig.1D**), and the posterior portion of the callosum was thinner (**Fig.1E**). These data suggest that TRIM67 is required for the midline-directed outgrowth and/or guidance of cortical axons, which form the corpus callosum.

**Fig. 1:**
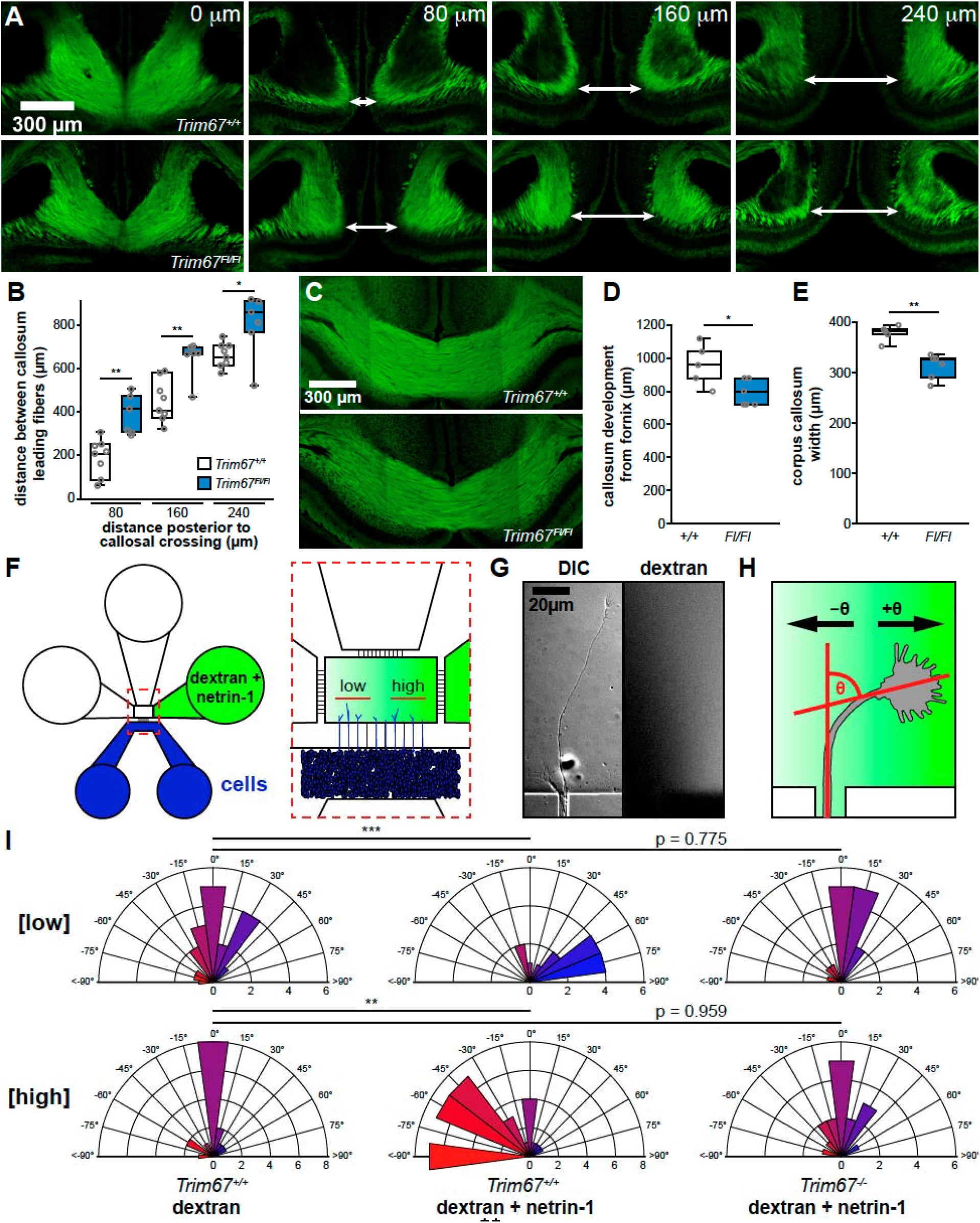
*Trim67* is required for axonal development and guidance *in vivo* and *in vitro*. **A)** Confocal micrographs of GFP in the corpus callosum of brains fixed immediately at postnatal day 0 (P0) from *Trim67^+/+^*:Tau-Lox-STOP-Lox-GFP:Nex-Cre and *Trim67^Fl/Fl,^*:Tau-Lox-STOP-Lox-GFP:Nex-Cre mice littermates. Arrows demarcate leading fibers of the corpus callosum in sections 80, 160 and 240 μm posterior to the final connection of the callosum. **B)** Individual data points and box and whisker plots of the distance between leading fibers of the corpus callosum at P0. **C)** Confocal micrographs of the corpus callosum at four days of age (P4) 160 μm anterior to the final connection of the callosum. Quantification of the extent of callosal development expressed as **D)** distance from the fornix to the separation of the callosal leading fibers and **E)** callosal width at the midline eight sections posterior to the fornix at P4. **F)** Schematic of microfluidic axon guidance chambers. **G)** Example axon extending from a microgroove into the axon guidance chamber, and fluorescent dextran used to visualize the gradient. **H)** Diagram depicting axon turning angle measured between a line bisecting the axonal growth cone and a line overlapping and parallel to the axon as it exits microgroove. **I)** Rose plots of embryonic cortical neuron axon turning angles in a gradient of fluorescent dextran or dextran + netrin-1. Low concentration denotes the four microgrooves furthest from the gradient source while the four microgrooves closest to the source are denoted high concentration (Taylor et al., 2015). Positive turning angles represent axon turning toward the netrin-1 source; negative angles represent axon turning away from the source. * - p < 0.05, ** - p < 0.01, *** - p < 0.005.

Since loss of murine *Dcc* or the gene encoding netrin-1 (*Ntn1*) result in agenesis of the corpus callosum (Fazeli et al., 1997; Marsh et al., 2017), and our previous work revealed that TRIM67 interacts with DCC (Boyer et al., 2018), we hypothesized the TRIM67-dependent defects in the corpus callosum could arise from axon guidance failures. To investigate the role of TRIM67 in netrin-dependent axon turning, we employed microfluidic axon guidance chambers to establish a stable gradient of netrin-1 (Taylor et al., 2015) and measured axon turning angles of cortical neurons cultured from *Trim67^+/+^* and *Trim67^-/-^* embryos (**Fig.1F-H**). As previously reported, in *Trim67^+/+^* cells, we observed positive turning angles indicative of attractive turning in a low concentration range of the netrin-1 gradient (approximately 40-220ng/mL), but not in a dextran-only gradient (**Fig.1I**). At the higher end of the gradient (approximately 550-600ng/mL), *Trim67^+/+^* axons exhibited a negative turning angle, indicative of repulsion from netrin-1. However, axons of *Trim67^-/-^* neurons did not show a turning response to netrin-1 in either range, similar to *Trim67^+/+^* cells in the dextran gradient (**Fig.1I**). These results demonstrate a requirement for TRIM67 in netrin-1 dependent axon turning *in vitro*, consistent with *in vivo* defects in the formation of the netrin sensitive corpus callosum.

### Netrin increases TRIM67 localization to filopodia tips

TRIM67 is enriched in the developing cortex and present in axonal growth cones (Boyer et al., 2018). We next characterized the netrin sensitivity of TRIM67 localization in the growth cone. TRIM67 exhibited a punctate pattern of localization in axonal growth cones, specifically in the core region, the edge of the lamellipodium, and the distal 0.5 μm of filopodia tips (**Fig.2A-B**). A similar tip localization is apparent for myc-tagged TRIM67 transfected into embryonic cortical neurons and imaged by structured illumination microscopy (**Fig.S1A**). This is reminiscent of the localization of filopodial tip complex proteins such as TRIM9 (Menon et al., 2015), VASP (Lebrand et al., 2004), and DCC (Shekarabi and Kennedy, 2002). Intriguingly acute netrin-1 treatment enhanced the filopodial tip localization of TRIM67 (**Fig.2C-E**, p = 0.0354), suggesting that TRIM67 is located closer to the tip of filopodia following netrin treatment. This supports the hypothesis that TRIM67 is poised to modulate netrin-1 dependent axonal responses.

**Fig. 2:**
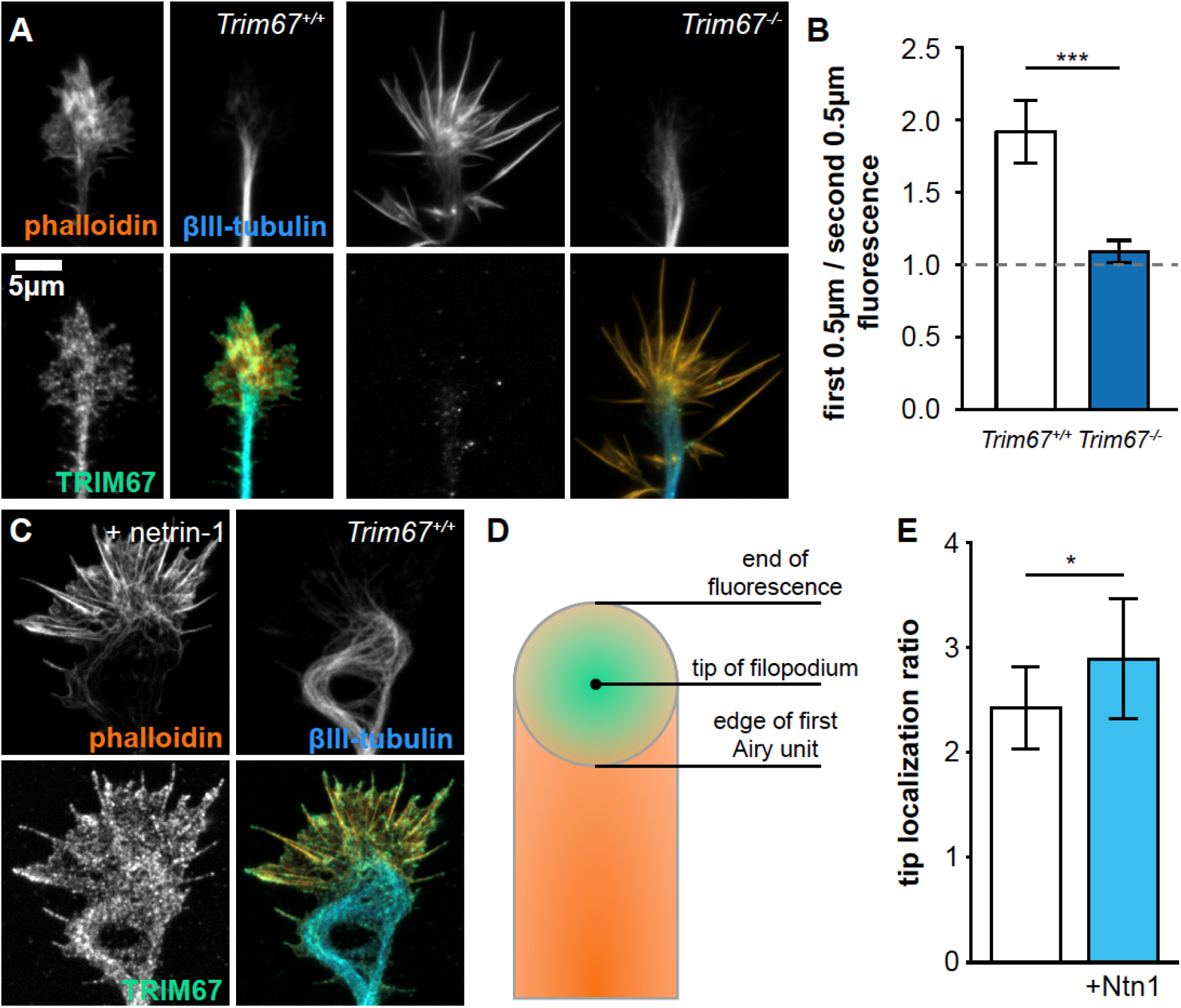
TRIM67 localization to filopodia tips is enhanced by netrin-1. **A)** Immunocytochemistry (ICC) of filamentous actin (phalloidin), ß-III-tubulin, and TRIM67 in axonal growth cones of primary neurons isolated from *Trim67^+/+^* and *Trim67^-/-^* embryonic cortices. **B)** TRIM67 fluorescence in the first 0.5 μm from the tip of the filopodium to the next 0.5 μm. **C)** ICC of an axonal growth cone from a *Trim67^+/+^* cortex treated with netrin-1. **D)** Diagram showing the Airy disk of a fluorescent protein at the tip of a filopodium (green) and fluorescence of a protein along the filopodium (orange). **E)** Tip proximity of TRIM67 in filopodia, quantified as the fluorescence ratio of the center to the edge of the first Airy unit (AU). Error bars denote SEM. * - p < 0.05, *** - p < 0.005.

### Axonal netrin-1 responses require TRIM67

We exploited scanning electron microscopy of cultured neurons to acquire a three dimensional (3D) gross overview of growth cone responses to netrin, and potential defects in *Trim67^-/-^* neurons (**Fig.3A**). We hierarchically categorized the 3D morphology of the growth cones as either flat (fully apposed to coverslip), superseded by curled (peripheral structures away from coverslip), then dorsal (possessing lamellipodial ruffles or filopodia on the dorsal surface), and finally nonadherent (growth cone fully separated from coverslip, **Fig.S1B**). Netrin-1 treatment promoted non-flat growth cone morphologies in *Trim67^+/+^* neurons (**Fig.S1C**, p = 0.040 by Fisher’s exact test). The growth cones of *Trim67^-/-^* axons were shifted towards nonflat morphologies (p = 0.011), and this distribution was not affected by netrin-1 (p = 0.310).

**Fig. 3:**
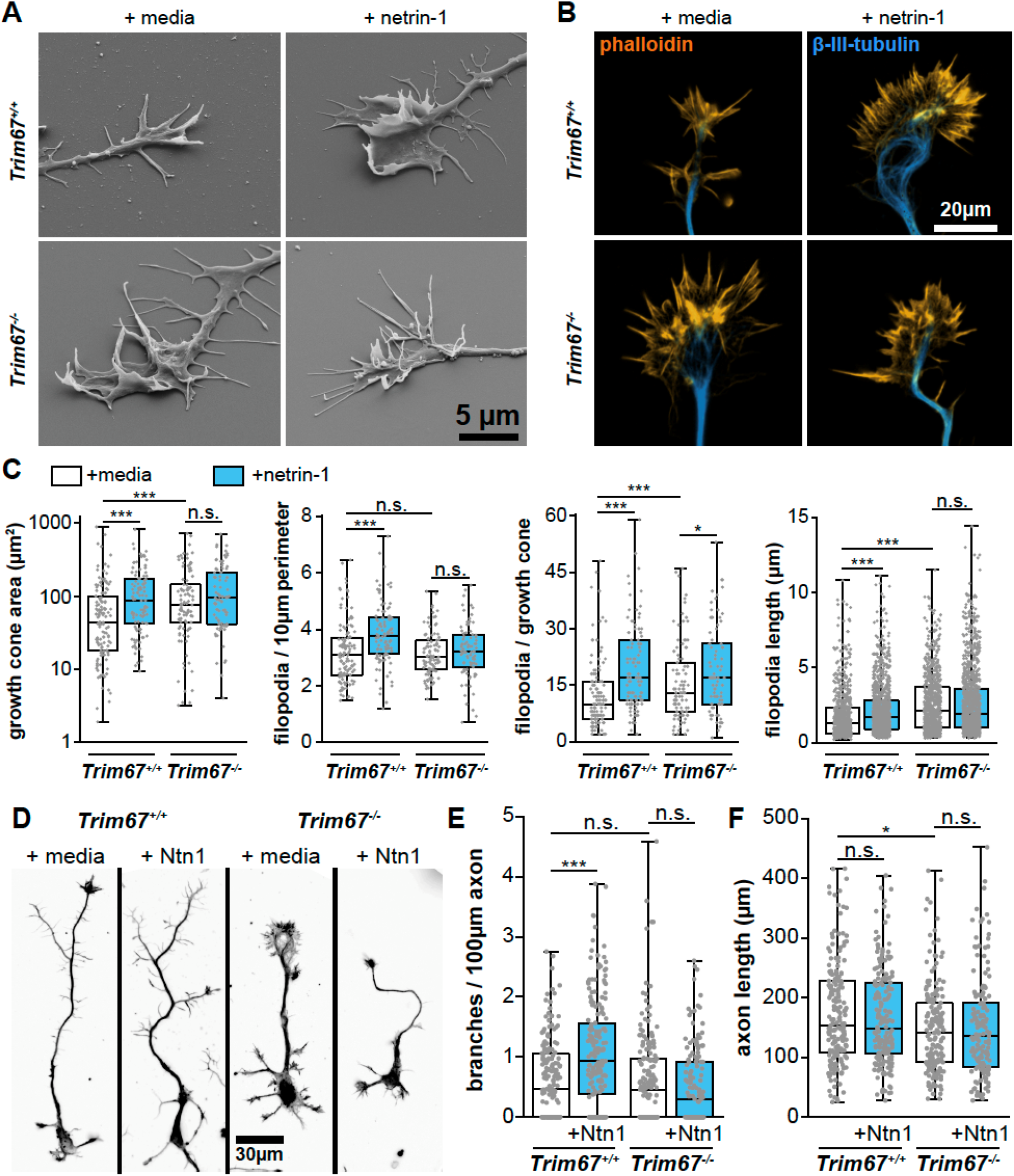
TRIM67 is required for axon and growth cone responses to netrin-1. **A)** Scanning electron micrographs of axonal growth cones from embryonic *Trim67^+/+^* and *Trim67^-/-^* cortices treated with sham media or media containing netrin-1. **B)** Growth cones from primary neuronal cultures stained for filamentous actin (phalloidin), β-III-tubulin, and TRIM67. **C)** Quantification of growth cone responses to netrin-1, including increase in growth cone area, filopodial density, filopodia number, and filopodia length. **D)** Micrographs of neurons cultured for 3 days *in vitro* including a final 24 hours with addition of media or netrin-1, shown as the combined fluorescence of staining for both filamentous actin (phalloidin) and β-III-tubulin. E) Quantification of axon branching per 100 μm axon length and of **F)** total axon length. * - p < 0.05, *** - p < 0.005, n.s. - p > 0.05.

The non-responsiveness of *Trim67^-/-^* growth cone morphology to netrin-1 prompted a thorough assessment of growth cone responses to netrin, including size, filopodial number, and filopodial length (**Fig.3B-C**). As reported (Menon et al., 2015), netrin-1 treatment increased growth cone area (p = 5.34E-4), filopodia density (p = 7.08E-7), filopodia number (p = 1.64E-7), and filopodial length (p = 7.97E-12) in *Trim67^+/+^* growth cones (**Fig.3C**). However, netrin-1 treatment had no effect on these parameters in *Trim67^-/-^* growth cones (area p = 0.896, filopodial density p = 0.434, filopodial number p = 0.2176, filopodial length, p = 0.808). This indicates TRIM67 is required for growth cone responses to netrin-1. Basally, filopodia of *Trim67^-/-^* growth cones were longer (p = 4.27E-23), and growth cones were larger (p = 0.00695) than wild-type counterparts. We investigated whether TRIM67 was similarly required for responses to other guidance cues (**Fig.S1D**). As reported previously (Szebenyi et al., 2001) treatment with FGF increased growth cone area and number of filopodia (**Fig.S1E**, area, p = 8.13E-5; filopodia, p = 2.38E-7). These effects were also observed in *Trim67^-/-^* neurons (**Fig.S1E**, area, p = 0.007; filopodia, p = 0.00722). Slit2N treatment caused a collapse of growth cones, as indicated by a decrease in the growth cone area in both *Trim67^+/+^* (p = 0.00291) and *Trim67^-/-^* (p = 0.00101) neurons (**Fig.S1E**).

Netrin also promotes axon branching in cortical neurons (Dent et al., 2004; Winkle et al., 2014). We assessed whether TRIM67 was required for axon branching after a 24 hour addition of netrin (**Fig.3D,E**). Netrin-1 increased the density of branches along *Trim67^+/+^* axons (p = 9.43E-7). In *Trim67^-/-^* axons this branching response was absent (p = 0.543), and there was no change in the branch density of untreated axons (p = 0.423). Basally *Trim67^-/-^* axons were shorter than their wild-type counterparts (p = 0.0133), though there was no effect on axon length from netrin-1 treatment in either genotype (**Fig.3F**, *Trim67^+/+^*, p = 0.711; *Trim67^-/-^*, p = 0.858). To determine whether the regulation of axon branching by TRIM67 was specific to netrin-1, we treated neuron cultures for with FGF or Slit2N (**Fig.S1F**). As reported (Szebenyi et al., 2001; Wang et al., 1999), both morphogens increased axon branching in *Trim67^+/+^* cortical neurons (**Fig.S1G**; FGF, p = 2.78E-4; Slit2N, p = 0.00175). Axons of *Trim67^-/-^* neurons were also more branched following treatment with FGF (p = 7.43E-4) or Slit2N (p = 0.00173). Along with the results of growth cone analysis, these data suggest that TRIM67 is not required for axonal responses to all guidance cues, but may be specific to netrin-1.

### Functional analysis of TRIM67 protein domains

After establishing the necessity for TRIM67 in axonal responses to netrin-1, we performed rescue experiments using full length TRIM67 or domain mutants of TRIM67 (**Fig.S2A**). The localization of most constructs was qualitatively similar to the full-length protein (**Fig.S2B**); however, qualitatively the ΔCC and Nterm constructs appeared more diffuse. Only full-length TRIM67 rescued the increase in growth cone area in response to netrin-1 treatment (**Fig.S2C**, p = 2.40E-5). However, all constructs containing the COS and FN3 domains rescued basal growth cone area, except for the ligase dead variant (**Fig.S2C**). Full-length TRIM67 also rescued the increase in filopodial density following netrin-1 treatment (**Fig.S2D**, p = 5.36E-7). Intriguingly, the only other construct which rescued filopodial density was the Nterm (p = 2.65E-5), suggesting that the N-terminus is sufficient for netrin sensitive filopodia-associated functions of TRIM67, but that a partial C-terminus may inhibit these functions. Most constructs did reduce the basal filopodial length, but not necessarily the netrin-dependent increase in length (**Fig.S2E**). These data suggest that all domains of TRIM67, as well as ligase activity are necessary to fully rescue all growth cone responses to netrin-1. Additionally, the COS and FN3 domains are required for TRIM67 to constrain the size of the growth cone lamellipodium in the absence of netrin.

We next explored the function of these TRIM67 domain mutants in netrin-1 dependent axon branching (**Fig.S3A**). Axon branching in response to netrin-1 was only rescued by full-length TRIM67 (**Fig.S3B**, p = 9.93E-7). Thus, the axonal defects in response to netrin are caused by loss of *Trim67*. These results suggest that like growth cone regulation, all domains of TRIM67 are necessary for proper netrin-dependent axon branching. Intriguingly, expression of TRIM67ΔCOS introduced a suppression of branching by treatment with netrin-1 (p = 0.00713). We conclude that TRIM67-dependent regulation of netrin responses is likely complex, as it requires each of the domains of TRIM67, and thus potentially distinct functions of the protein.

### Filopodia dynamics are regulated by TRIM67

We next assessed the dynamics of axonal growth cone filopodia using time-lapse microscopy (**Fig.4A, MovieS1**). In agreement with a previous study (Menon et al., 2015), we found that 40 min treatment with netrin-1 increased the lifetime of filopodia in *Trim67^+/+^* neurons (**Fig.4B**). Filopodia lifetime was basally longer in *Trim67^-/-^* growth cones and was reduced by addition of netrin-1. This decrease in lifetime could be attributed in part to an increase in the lateral buckling or folding of filopodia following netrin-1 treatment in *Trim67^-/-^* neurons (**Fig.4C**). We found that both the protrusion and retraction speed of the tips of filopodia were higher in *Trim67^-/-^* growth cones than in *Trim67^+/+^* growth cones, but that there was no effect on these rates with netrin-1 in either genotype (**Fig.4D**). However, the duration of individual filopodial retraction events was shorter in *Trim67^+/+^* growth cones following netrin-1 treatment, and this effect was absent in *Trim67^-/-^* filopodia (**Fig.4E**). Together, these data suggest that TRIM67 is required for filopodial growth dynamics to respond properly to netrin-1.

**Fig. 4:**
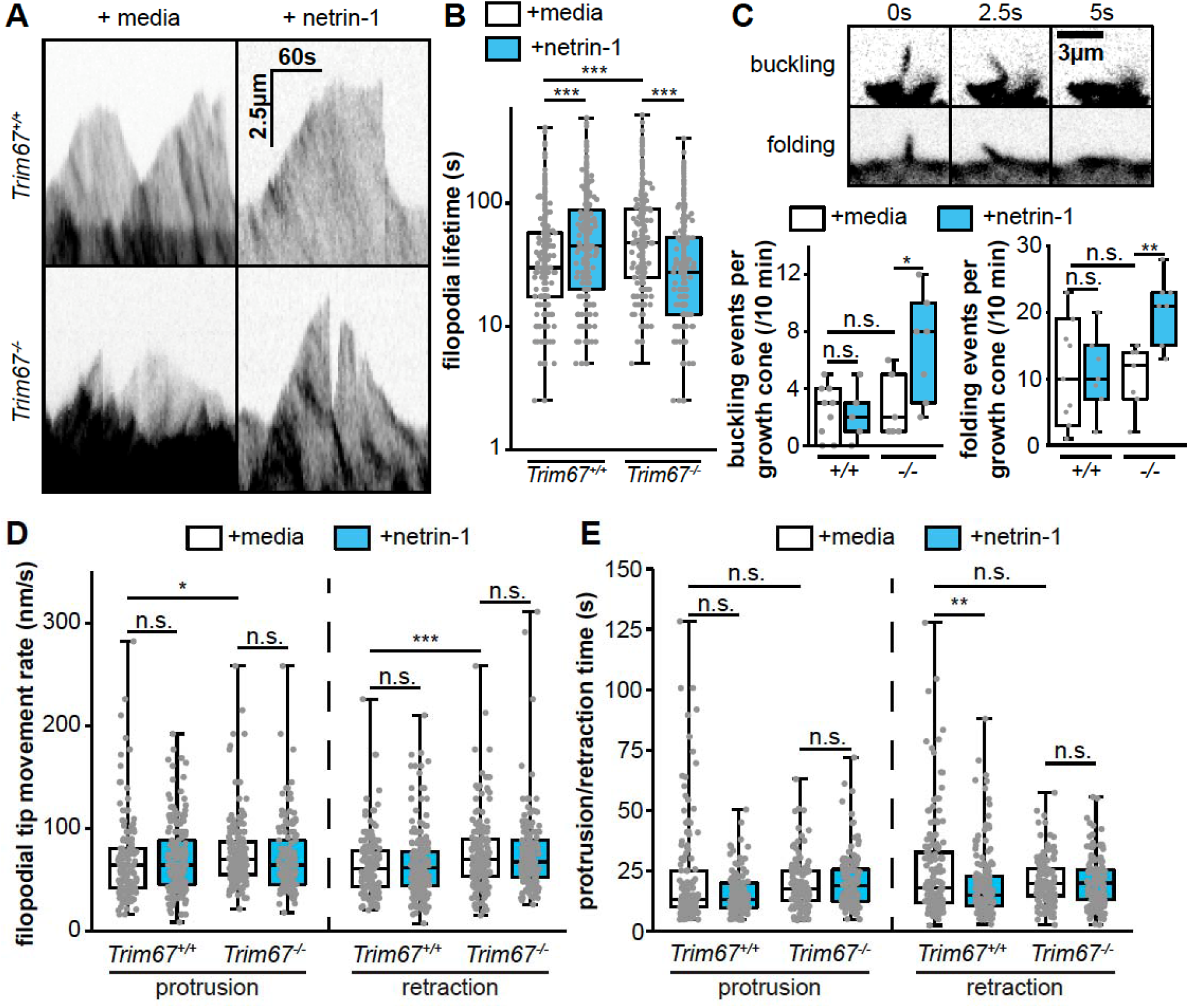
TRIM67 regulates filopodial growth and dynamics. **A)** Kymographs of filopodia from cultured primary embryonic cortical neurons expressing mcherry. **B)** Quantification of filopodial lifetime and **C)** both filopodial buckling and folding events during the course of 10 min time-lapse of axonal growth cones. **D)** Rate of filopodial tip movement and **E)** duration of individual filopodial protrusion and retraction periods following media sham or netrin treatment. * - p < 0.05, ** - p < 0.01, ***, p < 0.005, n.s. − p > 0.05.

### TRIM67 interacts and localizes with the filopodial actin polymerase VASP

Previous work demonstrated the Ena/VASP family of actin polymerases are required for filopodial responses to netrin-1 (Lebrand et al., 2004), and that VASP is regulated by TRIM9-dependent ubiquitination to mediate netrin-1 filopodial response (Menon et al., 2015). Since TRIM9 and TRIM67 are highly similar and interact (Boyer et al., 2018), we hypothesized that TRIM67 may also interact with VASP. Indeed, immunoprecipitation of Myc-TRIM67 or a mutant lacking the RING domain (MycTRIM67ΔRING) from HEK293 cells co-precipitated GFP-VASP (**Fig.5A**). The TRIM67:VASP complex was maintained in *TRIM9^-/-^* HEK293 cells, indicating the TRIM67:VASP interaction occurs independently of TRIM9 (**Fig.5A**). To map the domains of TRIM67 necessary for VASP interaction, we we generated a HEK293 cell line in which *TRIM67* was deleted via CRISPR/Cas9 genome editing (*TRIM67^-/-^* HEK293, **Fig.S4**) and performed co-immunoprecipitation assays using domain-deletion constructs of TRIM67. The coiled-coil domain of TRIM67 was required for co-immunoprecipitation of VASP, while ligase function was not (**Fig.5B**). Since interacting proteins often colocalize, we investigated whether tagRFPt-tagged TRIM67 and GFP-VASP colocalized in neurons by live TIRF microscopy (**Fig.5C**). Colocalization was quantified at growth cone filopodia tips and was higher than Fay-randomized controls (**Fig.5D**), indicating TRIM67 and VASP colocalize at filopodia tips. Consistent with co-immunoprecipitation results suggesting that the coiled-coil domain of TRIM67 is required for interaction with VASP; tagRFPt-TRIM67ΔCC showed decreased colocalization with GFP-VASP at filopodial tips. By coimmunoprecipitation we also found an interaction between TRIM67 and Ena/VASP family members Mena (**Fig.S4F**) and EVL (**Fig.S4G**).

**Fig. 5:**
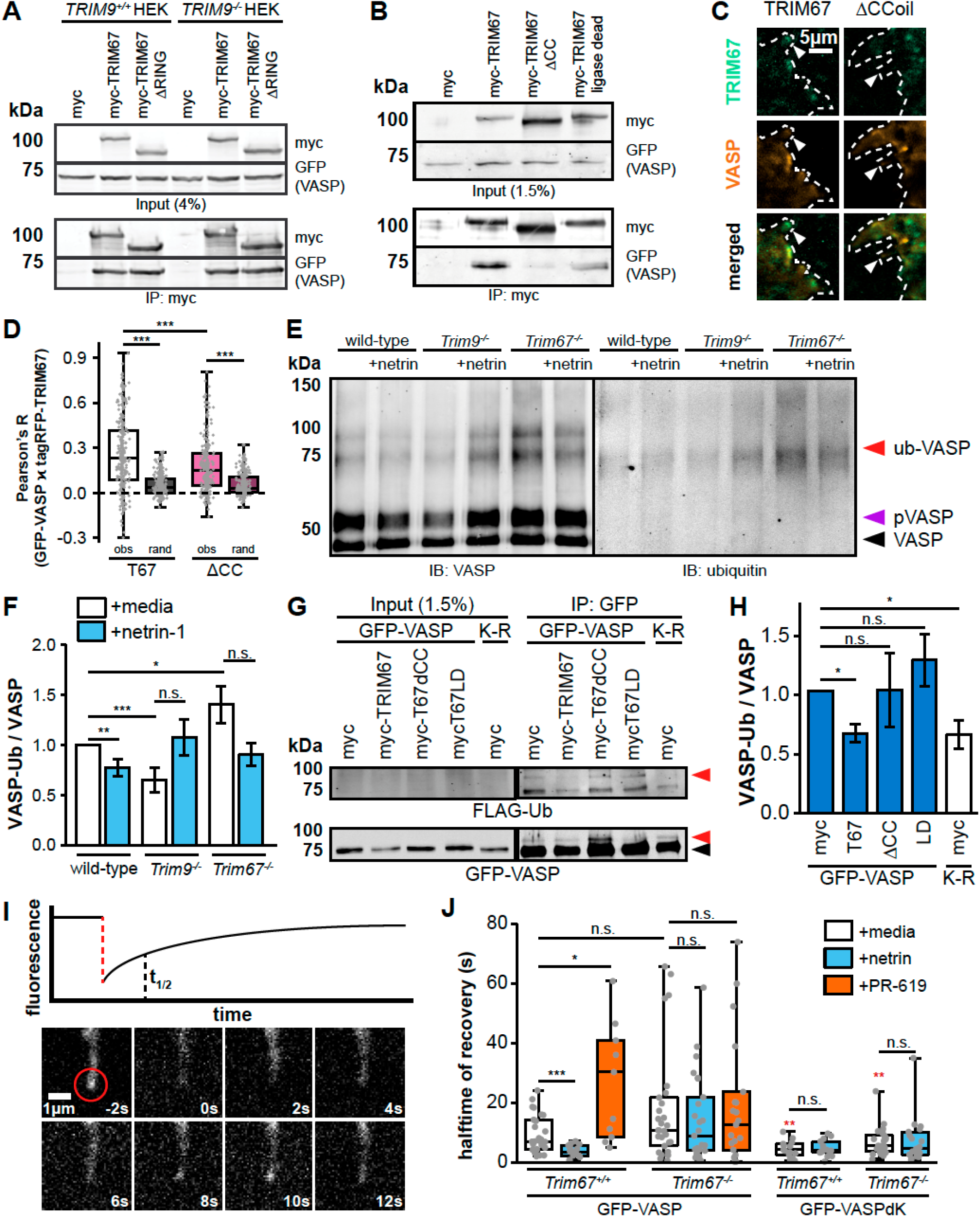
TRIM67 inhibits the ubiquitination of the actin polymerase VASP. **A)** Coimmunoprecipitation assays from *TRIM9^+/+^* or *TRIM9^-/-^* HEK293 cells transfected with GFP-VASP and myc or myc-tagged TRIM67 constructs demonstrate an interaction between TRIM67 and VASP that is independent of TRIM9. **B)** Coimmunoprecipitation assays from HEK293 cells transfected with the shown TRIM67 and VASP constructs, showing requirement for the TRIM67 coiled-coil domain for the TRIM67:VASP interaction. **C)** Colocalization between GFP-VASP and tagRFPt-tagged constructs of TRIM67 in murine embryonic cortical neurons. **D)** Quantification of the colocalization (□) between VASP and TRIM67 (R_(obs)_) in filopodia compared to Fay-randomized controls (R_(rand)_). **E)** Western blot of VASP immunoprecipitated from denatured cultured embryonic cortical lysate showing a band which co-migrates and co-labels with ubiquitin (VASP-Ub, red arrowhead), roughly 24kDa heavier than unmodified VASP, and a phosphorylated VaSp (pVASP) band roughly 2 kDa heavier than unmodified (purple arrowhead). **F)** Quantification of VASP-Ub relative to total VASP levels, normalized to untreated wild-type of each experiment. Bars are averages of 5-7 experiments ± SEM. **G)** Similar ubiquitination-precipitation assays of GFP-VASP expressed in HEK293T cells lacking *TRIM67* expressing indicated myc-TRIM67 constructs, along with FLAG-ubiquitin. A VASP band that comigrates with FLAG-Ub appears approximately 24kDa heavier than unmodified VASP (red arrowhead). **H)** Quantification of VASP-Ub, quantified from FLAG signal relative to total GFP-VASP, normalized to the myc control condition. **I)** Diagram of a fluorescence recovery curve following photobleaching of GFP-VASP at the tip of a filopodium with a representation of the halftime of recovery (t_1/2_), alongside an image montage of the FRAP of GFP-VASP in a transfected embryonic cortical neuron. **J)** Quantification of the FRAP t_1/2_ of GFP-VASP or GFP-VASP^K-R^ in embryonic cortical neurons treated with netrin-1 or the deubiquitinase inhibitor PR-619. Statistical comparisons in red are with respect to the GFP-VASP FRAP t_1/2_ in untreated cells of the same genotype. * - p < 0.05, ** - p < 0.01, *** - p < 0.005, n.s. − p > 0.05.

### TRIM67 antagonizes VASP ubiquitination

We previously reported that TRIM9-dependent non-degradative ubiquitination of VASP negatively impacted VASP dynamics and filopodial stability, all of which were reversed by netrin (Menon et al., 2015). This work detected no ubiquitination of other Ena/VASP family members Mena and Evl, and similar results were found here with Mena (**Fig.S4H**). Due to the structural similarities between TRIM9 and TRIM67 and their conserved coiled-coil mediated interaction with VASP, we investigated how TRIM67 modulated VASP ubiquitination. Consistent with previous work, we found basal level of VASP ubiquitination in *Trim67^+/+^* neurons, which was decreased in response to netrin-1 treatment (**Fig.5E,F**, p = 0.00805). Further we confirmed previous results that VASP ubiquitination was decreased in the absence of *Trim9* (p = 0.00220). Surprisingly, in *Trim67^-/-^* neurons there was an increase in VASP ubiquitination and a trend towards decreased VASP ubiquitination following netrin-1 treatment (*Trim67^+/+^* vs. *Trim67^-/-^* p = 0.018, netrin effect p = 0.0573). Again, we conclude this ubiquitination is likely not associated with degradation of VASP, as changes in levels of VASP protein were not detected in *Trim67^-/-^* or *Trim9^-/-^* brain lysates (**Fig.S4I-J**). We performed similar ubiquitination assays of GFP-VASP in HEK293 cells and found high levels of VASP ubiquitination in the absence of TRIM67 (**Fig.5G,H**). Introduction of myc-tagged TRIM67 decreased VASP ubiquitination (p = 0.0300), consistent with endogenous VASP ubiquitination in neurons. The TRIM67-dependent inhibition of VASP ubiquitination required the coiled-coil domain of TRIM67 as well as ligase function, as neither mutant decreased ubiquitination compared to myc-transfected *TRIM67^-/-^* HEK cells (**Fig.5G,H**, ΔCC, p = 0.874; LD, p = 0.656). This suggests that the interaction of TRIM67 with VASP or another protein as well as TRIM67 E3 ligase function, are necessary for inhibiting VASP ubiquitination. To confirm the molecular weight shift of VASP and co-migration of ubiquitin were indicative of increased ubiquitination in the absence of TRIM67, we exploited a construct of GFP-VASP harboring nine lysine residues mutated to arginine (VASP^K-R^). These mutations were previously shown to decrease VASP ubiquitination (Menon et al., 2015). Similarly here VASP^K-R^ was not ubiquitinated in the absence of TRIM67 (**Fig.5G,H**, 0.0150).

### VASP ubiquitination slows VASP dynamics at filopodia tips

Using fluorescence recovery after photobleaching (FRAP) of GFP-VASP, we previously found that when VASP was ubiquitinated, the fluorescence recovery halftime (t_1/2_) was slow, whereas when VASP was not ubiquitinated the FRAP t_1/2_ was fast (Menon et al., 2015). We therefore performed FRAP assays in embryonic cortical neurons transfected with GFP-VASP (**Fig.5I**) to determine if loss of *Trim67* also altered VASP dynamics at filopodia tips. Consistent with our previous work, treatment with netrin-1 caused a reduction in the t_1/2_ of filopodial GFP-VASP (**Fig.5J**), indicating more rapid dynamics of VASP at filopodial tips when VASP ubiquitination was reduced (Menon et al., 2015). In *Trim67^-/-^* neurons there was no change in t_1/2_ with netrin treatment, matching the pattern found in our *in vitro* ubiquitination assays. To test whether this effect on FRAP t_1/2_ was due to ubiquitination, we performed FRAP experiments using GFP-VASP^K-R^. In both *Trim67^+/+^* and *Trim67^-/-^* neurons the FRAP t_1/2_ of GFP-VASP^K-R^ was lower than that of GFP-VASP, and there was no effect of netrin on the t_1/2_ in either genotype. To assay effects of increased VASP ubiquitination on FRAP t_1/2_ we treated neurons with PR-619; in *Trim67^+/+^* neurons PR-619 increased GFP-VaSp FRAP t_1/2_, consistent with an increase in ubiquitination (**Fig.5J**). However, the VASP t_1/2_ was not affected in *Trim67^-/-^* neurons treated with PR-619, suggesting that deubiquitinase inhibition does not impact the mobility of VASP in the absence of *Trim67* as inhibition does in wild-type cells. We see no difference in the percent of VASP which recovers after photobleaching between genotypes or treatment conditions (**Fig.S5A**). Together these data suggest that VASP ubiquitination correlates with the rate of FRAP recovery, as previous work suggested (Menon et al., 2015), and that TRIM67 is required for the proper netrin-1 dependent increase in VASP mobility at filopodia tips.

### TRIM67 negatively regulates TRIM9 through competition for binding with substrates

Our data are consistent with the hypothesis that TRIM67 antagonizes the TRIM9-dependent ubiquitination of VASP, and previous work showed that TRIM67 and TRIM9 interact (Boyer et al., 2018). Therefore, we investigated whether TRIM67 and TRIM9 colocalize in cultured neurons, such that TRIM67 may inhibit TRIM9-dependent ubiquitination of VASP. In *Trim9^-/-^:Trim67^-/-^* embryonic cortical neurons GFP-TRIM9 and tagRFPt-TRIM67 colocalized significantly when compared to Fay-randomized controls (**Fig.6A,B**). Netrin treatment did not detectably alter the colocalization of these two proteins.

**Fig. 6:**
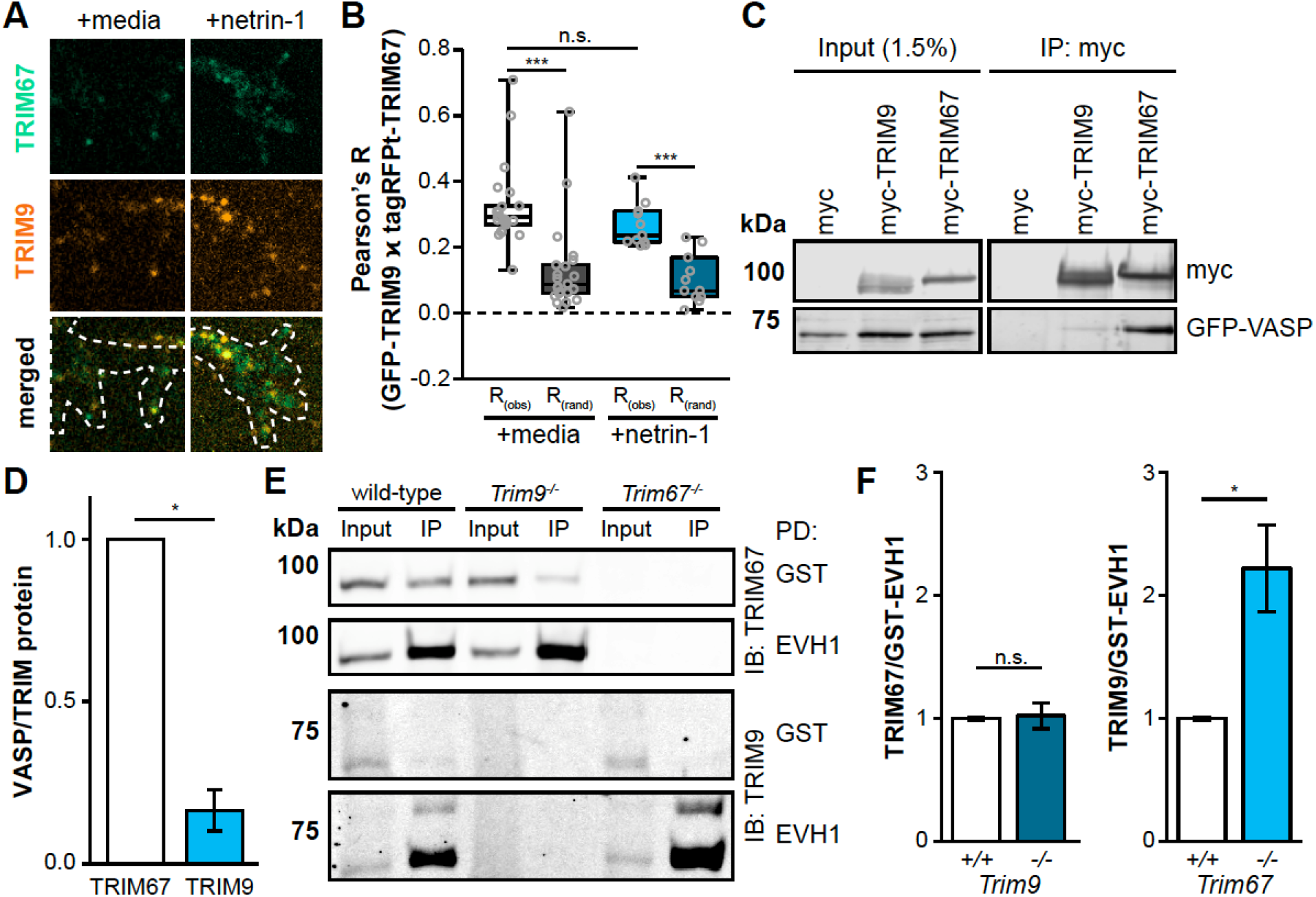
TRIM67 competitively inhibits the TRIM9 interaction with VASP. **A)** Colocalization of GFP-TRIM9 and tagRFPt-tagged TRIM67 in embryonic cortical neurons. B) Quantification of colocalization (□) between TRIM9 and TRIM67 (R_(obs)_) compared to Fay-randomized controls (R_(rand)_). **C)** Coimmunoprecipitations of myc-tagged TRIM9 or TRIM67 from HEK293 cells showing precipitated GFP-VASP. **D)** Quantification of co-precipitated GFP-VASP relative to myc-tagged TRIM protein normalized to the VASP/TRIM67 ratio of each experiment. **E)** Pulldowns from embryonic mouse cortical lysate using either GST or GST-EVH1 domain of VASP, probed for endogenous TRIM67 and TRIM9. F) Quantification of TRIM proteins precipitated by GST-EVH1 from lysates of indicated genotypes, normalized to wild-type levels. * - p < 0.05, *** - p < 0.005, n.s. − p > 0.05.

We hypothesized that TRIM67 might compete with TRIM9 for an interaction with VASP, as deletion of *Trim67* did not affect overall TRIM9 protein levels (**Fig.S4K,L**). Coimmunoprecipitation assays suggested a stronger interaction between TRIM67 and VASP than between TRIM9 and VASP (**Fig.6D,E**, p = 0.0211). The Ena/VASP homology domain 1 (EVH1) of VASP, which interacts with TRIM9 (Menon et al., 2015), precipitated both TRIM67 and TRIM9 from cortical lysate when tagged with glutathione S-transferase (GST), indicating both proteins interact with the same domain of VASP (**Fig.6F**). GST-EVH1 enriched indistinguishable amounts of endogenous TRIM67 from wildtype and *Trim9^-/-^* cortical lysate (**Fig.6F,G**, p = 0.878), indicating TRIM9 did not impair the TRIM67:EVH1 interaction. However, GST-EVH1 precipitated approximately two fold more TRIM9 in the absence of *Trim67* (**Fig.6F,G**, p = 0.0211), suggesting that TRIM67 competes with the interaction between TRIM9 and VASP.

In light of this competitive interaction, we hypothesized that TRIM67 functioned upstream of TRIM9, and that would predict that VASP ubiquitination and filopodial responses to netrin-1 in *Trim9^-/-^:Trim67^-/-^* cortical neurons would resemble *Trim9^-/-^* neurons (Menon et al., 2015). Indeed in *Trim9^-/-^:Trim67^-/-^* neurons we observed a decrease in VASP ubiquitination (**Fig.7A,B**, p = 0.00210), similar to in *Trim9 ^-/-^* neurons. Consistent with the hypothesis that ubiquitination of VASP slows its dynamics at filopodia tips, the FRAP t_1/2_ of GFP-VASP expressed in 2xKO embryonic cortical neurons was lower than in untreated wild-type neurons, and displayed an increase following netrin treatment or addition of PR-619 (**Fig.7C**). As with previous FRAP assays we saw no differences in % recovery with any condition (**Fig.S5B**) Analysis of *Trim9^-/-^:Trim67^-/-^* axonal growth cones (**Fig.7D**) showed that similar to those of *Trim9^-/-^* neurons, basal filopodial number and filopodial density increased in the absence of both TRIM proteins and did not increase in response to netrin-1 treatment (**Fig.7E**).

**Fig. 7:**
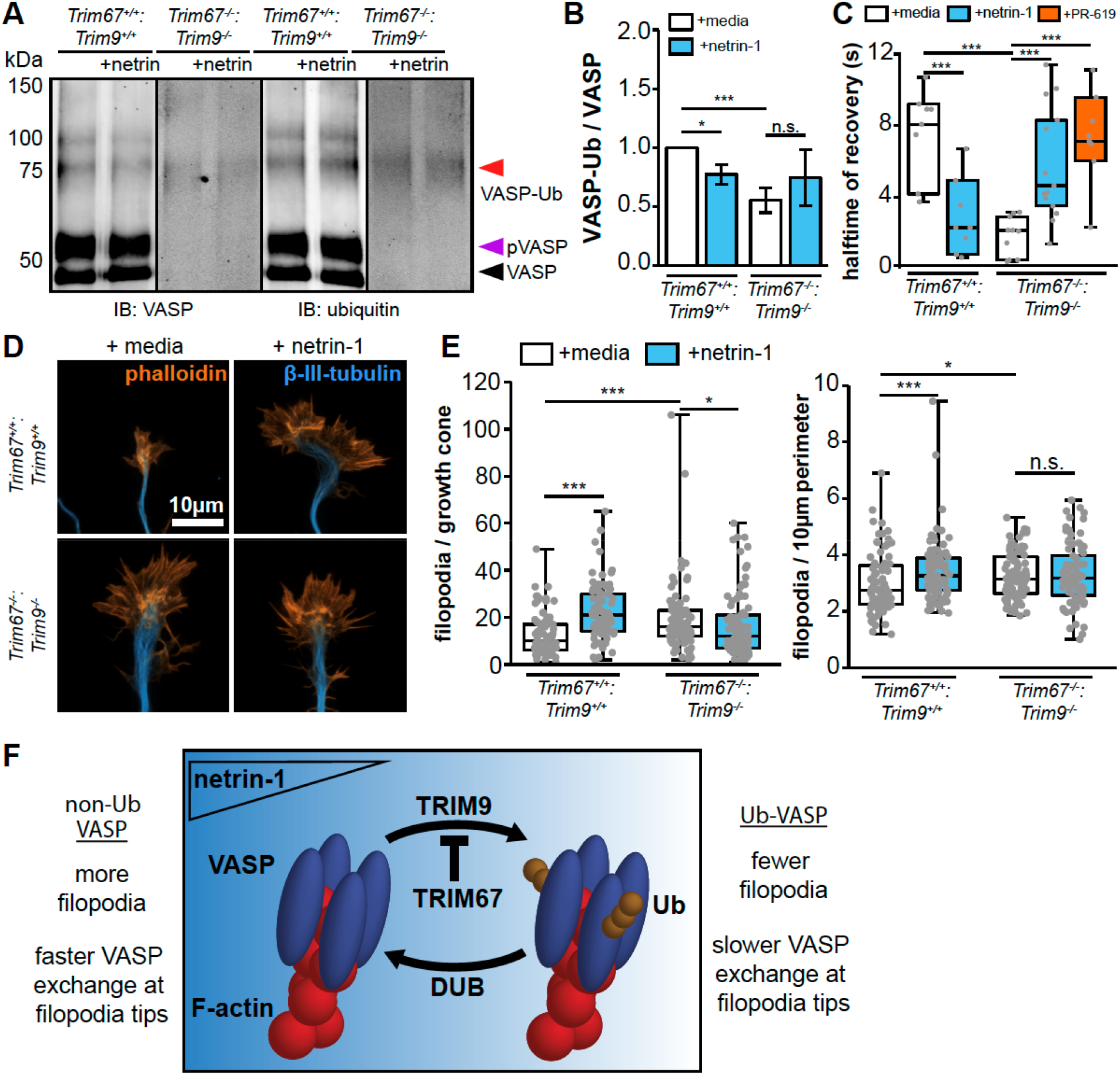
TRIM67 functions upstream of TRIM9 in the regulation of VASP and filopodia. **A)** VASP ubiquitination immunoprecipitation from *Trim9^-/-^:Trim67^-/-^* embryonic cortical neurons. **B)** Quantification of VaSp ubiquitination; bars are averages of 4-7 experiments ± SEM. C) FRAP halftime (t_1/2_) of GFP-VASP at filopodia tips in *Trim9^-/-^:Trim67^-/-^* embryonic cortical neurons treated with netrin-1 or PR-619. **D)** Growth cones of cortical neurons cultured from *Trim67^+/+^:Trim9^+/+^* or *Trim67^-/-^:Trim9^-/-^* embryos, treated with either netrin-1 or a media sham. **E)** Filopodia number per growth cone and numerical density in embryonic cortical neurons isolated from *Trim9^-/-^:Trim67^-/-^* brains. **F)** Model of TRIM67 in the regulation of VASP ubiquitination, and the role of VASP-Ub in the cell. * - p < 0.05, *** - p < 0.005, n.s. − p > 0.05.

## DISCUSSION

In this study we demonstrate that TRIM67 antagonizes the TRIM9-dependent ubiquitination of VASP and is crucial for axonal responses to netrin-1. This leads to our working model shown in **Fig.7F**, in which VASP is ubiquitinated by TRIM9, resulting in decreased stability of filopodia on axonal growth cones. This ubiquitination is antagonized by TRIM67, potentially via competition with TRIM9 for interaction with the EVH1 domain of VASP. We found that the ubiquitination and dynamics of VASP, the filopodial morphology, and netrin responses in a *Trim67:Trim9* double knockout neuron resemble that of a *Trim9^-/-^* neuron. This suggests that TRIM9 acts downstream of TRIM67. We hypothesize that VASP ubiquitination and the resultant short lived filopodia allow for efficient filopodial exploration of the extracellular environment. Consequently, when a filopodia encounters netrin, TRIM67 is recruited to filopodia tips, where it antagonizes VASP ubiquitination and increases filopodial lifetime, prior to axon turning.

### TRIM67 regulates brain development

These experiments support the hypothesis that TRIM67 is critical for aspects of brain development, particularly appropriate midline crossing of axons through the corpus callosum. In agreement with previous work showing that genetic deletion of *Trim67* results in thinning of the adult corpus callosum, we found a reduction in the midline-directed outgrowth of callosal fibers during development. This may be due to diminished attraction of *Trim67^-/-^* axons to netrin-1 at the midline, as we find that *Trim67^-/-^* axons are insensitive to netrin-1 *in vitro*. The reduced axon length observed in *Trim67^-/-^* cultured neurons may also contribute to this midline-crossing deficiency. Similar deficits in axon outgrowth and guidance in response to netrin-1 could contribute to other neuroanatomical defects seen in the brains of *Trim67^-/-^* mice. Additionally, our observation that TRIM67 is required for netrin-dependent axon branching *in vitro* may predict a reduction in axon arbor elaboration and subsequently innervation by netrin-1 sensitive neurons. These possibilities are intriguing, as several of the hypotrophies seen in *Trim67^-/-^* brains are in regions with evidence of netrin-1 secretion (Boyer et al., 2018; Fazeli et al., 1997; Serafini et al., 1996; Xu et al., 2010; Yung et al., 2015).

### TRIM proteins as models of paralog interference

Although TRIM67 may act as an E3 ubiquitin ligase (Yaguchi et al., 2012), our observations suggest it also functions as a competitive inhibitor to a closely related E3 ligase. Based on phylogenetic analysis (Boyer et al., 2018), the TRIM9 and TRIM67 paralog pair likely arose from a gene duplication event, producing two similar proteins in vertebrates. Duplication of gene products that form homodimers, as many TRIM proteins do (Koliopoulos et al., 2016; Sanchez et al., 2014), can lead to a phenomenon known as paralog interference, in which one member of the pair diverges and acts as an inhibitor for the other protein (Baker et al., 2013; Kaltenegger and Ober, 2015). While this behavior is usually seen in relation to transcription factors, the ability of TRIM67 to antagonize the interaction between TRIM9 and VASP and TRIM9-dependent VASP ubiquitination suggests that TRIM9 and TRIM67 may represent an example of paralog interference. This is supported by the observation that the ability of TRIM67 to antagonize TRIM9-dependent VASP ubiquitination in HEK293T cells required the domain of TRIM67 that mediates TRIM dimerization. Class 1 TRIM proteins in mammals are all paralog pairs (Short and Cox, 2006), and as such, the interaction shown here for TRIM9 and TRIM67 may be relevant for other class 1 TRIM pairs: TRIM1 and TRIM18, TRIM36 and TRIM46. Given the frequency of whole-genome duplication events throughout animal evolution (Glasauer and Neuhauss, 2014; Van De Peer et al., 2009; Robertson et al., 2017; Song et al., 2016) and the consequently conserved dimerization domains of RING domain-containing E3 ligases (Choo and Hagen, 2012; Liew et al., 2010; Metzger et al., 2014), competitive inhibition by paralogs may be a common regulatory mechanism. Future studies will be necessary to explore whether other TRIM proteins display similar inhibition in paralog pairs.

### TRIM67 is a protein with diverse functions

The results of our structure-function rescue assays suggest that additional functions, interaction partners, and substrates of TRIM67 remain to be identified. For example, the rescue of growth cone area was dependent upon the COS and FN3 domains of TRIM67, but did not appear to be correlate with the ubiquitination state of VASP. This suggests TRIM67-dependent regulation of other growth cone proteins regulates growth cone size. Likewise, the peculiar gain-of-function in neurons transfected with TRIM67ΔCOS, resulting in a suppression of branching by netrin-1 requires additional investigation. Other members of the class 1 TRIM proteins interact with microtubules via the COS domain (Cainarca et al., 1999; Cox, 2012; Wright et al., 2016, van Beuningen et al., 2015). Regulation of microtubules may be responsible for the branching phenotype and lamellipodial reorganization and size. This suggests another intriguing avenue for future research, and could facilitate the identification of substrates of TRIM67.

### The puzzle of VASP ubiquitination

Our findings here bolster the hypothesis that nondegradative ubiquitination of VASP alters or inhibits the functionality of the polymerase in growth cones, and that changes in the degree of ubiquitination of VASP are associated with netrin-1 dependent axonal responses. Previously we found that a cycle of VASP ubiquitination and deubiquitination was necessary for filopodial responses to netrin-1 (Menon et al., 2015). Building upon this, our data suggest that the TRIM9-dependent ubiquitination of VASP is inhibited by TRIM67. Therefore, we propose a coordination of VASP ubiquitination by the yin-and-yang-like pair of TRIM9 and TRIM67. This establishes new avenues of inquiry: how does ubiquitination modify VASP function? Are other signaling pathways and cytoskeletal associated proteins regulated in this manner? The requirement of ubiquitination of a relatively small amount of VASP for appropriate netrin responses suggests ubiquitination potently regulates VASP function. As VASP functions as a tetramer at actin barbed ends, this could be the result of one modified VASP monomer altering the function of the entire tetramer. Further studies need to investigate how ubiquitination of VASP alters its function on the molecular level or changes the state of the tetramer. Such work could have intriguing implications for the regulation of other actin associated proteins that function as multimers, such as formins, which recent work has suggested are also regulated by nondegradative ubiquitination (Angeles Juanes and Piatti, 2016).

## ACKNOWLEDGMENTS

The authors would like to thank Anthony Mangan for assistance with immunoblotting and Caroline Monkiewicz and Vong Thoong for mouse breeding, husbandry, and genotyping. Funding from the National Institutes of Health supported this research: including R01GM108970 (SLG) and F31NS096823 (NPB), as well as funding for UNC Neuroscience microscopy core facility P30 NS045892 and U54 HD079124.

## AUTHOR CONTRIBUTIONS

Conceptualization, N.P.B. and S.L.G.; Methodology, N.P.B. and S.L.G.; Investigation, N.P.B., L.E.M. and F.L.U.; Writing – Original Draft, N.P.B., F.L.U. and S.L.G.; Writing – Review & Editing, N.P.B., L.E.M., and S.L.G.; Funding Acquisition, N.P.B. and S.L.G.; Supervision, S.L.G.

## EXPERIMENTAL MODEL AND SUBJECT DETAILS

### Animals

All mouse lines were on a C57BL/6J background and bred at UNC with approval from the Institutional Animal Care and Use Committee. Timed pregnant females were obtained by placing male and female mice together overnight; the following day was designated as E0.5 if the female had a vaginal plug. Sexes of early postnatal mice used for corpus callosum development assays were not recorded, due to the inaccuracy of standard sexing procedures before postnatal day 8 (Greenham and Greenham, 1977). *Trim9^-/-^, Trim67^-/-^, Trim67^fl/fl^*, and Nex-Cre mice were described (Boyer et al., 2018; Goebbels et al., 2006; Menon et al., 2015; Winkle et al., 2014). Double TRIM knockout mice were generated by crossing *Trim9^-/-^* and *Trim67^-/-^* mice, and then crossing resultant heterozygotes.

### Cortical neuron culture

Embryonic day 15.5 (E15.5) mice were removed from pregnant dams by postmortem cesarean section, and dissociated cortical neuron cultures were prepared as described previously (Kwiatkowski et al., 2007). Cortices were micro-dissected and neurons were dissociated with 0.25% trypsin for 15 minutes at 37°C, followed by quenching of trypsin with neurobasal medium supplemented with 10% FBS and 0.5mM glutamine. After quenching, cortices were gently triturated 15x in quenching medium, counted by hemocytometer, and spun in a tabletop centrifuge at 800 rpm for 7 minutes at room temperature. Trypsin quenching medium was aspirated and cells were gently resuspended in culture media and plated on Poly-D-lysine (Sigma)-coated coverglass or tissue culture plastic in Neurobasal medium supplemented with B27 (Invitrogen). To assay growth cones and filopodia, 600 ng/ml recombinant netrin-1, 24 ng/ml recombinant FGF-2 (MBL International) or 400 ng/mL Slit2N (PeproTech) was bath applied after 48 hrs *in vitro* for 40 min, or 250ng/mL netrin-1, 10 ng/mL FGF-2, or 100 ng/mL Slit2N was bath applied for 24 hours after 48 hours *in vitro*, followed by fixation with 4% paraformaldehyde in PHEM buffer. Cells were permeabilized for 10 minutes in 0.1% TritonX-100, blocked for 30 minutes in 10% BSA, and stained with indicated primary antibodies for 1 hour at room temperature. Following three washes, species appropriate fluorescent secondary antibodies were added and allowed to incubate for 1 hour at room temperature. Following three washes, cells were mounted in a TRIS/glycerol/n-propyl-gallate based mounting media for imaging. Widefield epifluorescence images of pyramidal-shaped neurons were analyzed. Growth cone perimeter and area were measured using ImageJ. Filopodium length was measured from the filopodium tip to lamellipodial veil. Number of filopodia was counted per growth cone, and density is reported per 10 μm of growth cone perimeter. To assay filopodia dynamics, dynamic colocalization and FRAP, time-lapse imaging was performed with a stage top incubator that maintained humidity, 37°C and 5% CO_2_ (Tokai Hit).

### HEK293 cell culture

HEK293 cells (female) were obtained from Dr. Simon Rothenfusser (Klinikum der Universität München). HEK293 cells were cultured in DMEM medium supplemented with 10% fetal bovine serum and maintained at 5%CO_2_ / 37°C.

### Generation of *TRIM67^-/-^* HEK cells

The generation of HEK293 *TRIM67^-/-^* cells was performed using CRISPR/Cas9 gene editing with single-guide RNA targeting three sets of sequences in Exon 1 of the *TRIM67* gene in HEK293 cells:

Set 1: 5’-TCCTGCTTTCCCGGGGATCG-3′; 5’-GGCAGGTGCTGCTCACCGTC-3′
Set 2: 5’-GCTCCTGCTTTCCCGGGGAT-3′; 5’-CAGGTGCTGCTCACCGTC-3′
Set 3: 5’-CTCCTGCTTTCCCGGGGATC-3′; 5’-GGCAGGTGCTGCTCACCGTC-3′

1.3 x 10^5 HEK293 cells were seeded and transfected in a 24-well plate with 250ng of an expression plasmid coding for a CMV promoter-driven nickase-variant Cas9-D10A-puromycin cassette and 250ng of the mixed U6-driven single-guide RNAs using Lipofectamine 2000 (Invitrogen) as per manufacturer protocol. After 24h transfected cells were selected for by treatment with 1ug/mL puromycin and incubated for 72 hours. Remaining cells were diluted to 0.5 cells/100uL and single cells were seeded in 96 well plates. Growing clones were expanded. Genomic DNA was extracted and the target region of interest was amplified by PCR to detect size differences. Positive clones were screened by PCR (Primers: 5’ CCTTCTCTCGCCCCTCAATC 3’, and 5’ GTGCTCCAGAGAGGCAGC 3’) and sequenced. A knockout cell clone was identified harboring frameshift mutations in the exon 1 region and verified by Western blot protein analysis. HEK293 cells were maintained in DMEM media with glutamine (Invitrogen), supplemented with 10% fetal bovine serum (Hyclone). *TRIM9^-/-^* HEK293 cells were described previously (Menon et al., 2015).

### Plasmids, antibodies and reagents

Plasmids encoding human TRIM9 cDNA, mouse TRIM67 and GFP-VASP K-R mutant were described previously (Boyer et al., 2018; Menon et al., 2015; Winkle et al., 2014). The following plasmids were acquired: mCherry (Clonetech), FLAG-Ub (Dr. Ben Philpot, UNC-Chapel Hill), pmscv-eGFP-Mena, pmscv-eGFP-EVL, pQE-EVH1, pGEX-6P-1-Pro (Dr. Frank Gertler, MIT), peGFP-N1-VASP (Dr. Richard Cheney, UNC). TRIM67 CC domain sequence (aa332-369 of murine TRIM67) was cloned into the pGEX-6P-1plasmid. Myc-tagged TRIM67 domain deletion constructs were generated by partial PCR of myc-TRIM67; TRIM67ΔRING (aa159-aa768); TRIM67ΔCC (aa1-aa335, lysine, glutamate, aa369-aa768); TRIM67ΔCOS (aa1-aa369, lysine, glutamate, 6x glycine, aa494-aa768); TRIM67ΔFN3 (aa1-aa497, lysine, glutamate, aa579-aa768); TRIM67ΔSPRY (aa1-aa632); TRIM67 N-terminus (aa1-aa369). TRIM67 ligase dead construct was generated by quick-change site directed mutagenesis of cysteines 7 and 10 to alanine (TGC -> GCC).

Antibodies include: rabbit polyclonal against TRIM67 (Boyer et al., 2018), rabbit polyclonal antibody against VASP (Dr. Frank Gertler, MIT), rabbit polyclonal against VASP (sc-13975, SCBT), mouse monoclonal against TRIM9 (H00114088-M01, Abnova), mouse monoclonal against c-Myc (9E10, SCBT), mouse monoclonal against human ßIII Tubulin (TujI SCBT), rabbit polyclonal against FLAG tag (F7425, Sigma), rabbit polyclonal against GFP (A11122 Invitrogen), mouse monoclonal against GFP (75-131 UC Davis Neuromab), chicken polyclonal against GFP (GFP-1010, aves LABS, inc.), ubiquitin (sc-8017, SCBT) and GAPDH (sc-166545, SCBT). Fluorescent secondary antibodies and fluorescent phalloidin labeled with AlexaFluor 488, AlexaFluor 568, or AlexaFluor647 were from Invitrogen. Recombinant netrin-1 was concentrated from HEK293 cells (Kennedy et al., 1994; Lebrand et al., 2004). PR-619 ((2,6-diaminopyridine-3,5-bis(thiocyanate), abcam), MG132 (81-5-15, American Peptide Company), Rhodamine B isothiocyanate–Dextran (Sigma-Aldrich, R9379).

### Transfection procedures

For transfection of plasmids, neurons were resuspended after dissociation in Lonza Nucleofector solution (VPG-1001) and electroporated with an Amaxa Nucleofector according to manufacturer protocol. HEK cells were transfected using Lipofectamine 2000 (Invitrogen) as per manufacturer protocol.

### Immunoblotting and precipitation assays

For ubiquitination assay MG132 and netrin-1 treated cells were lysed in IP buffer (20 mM Tris-Cl, 250 mM NaCl, 3 mM EDTA, 3 mM EGTA, 0.5% NP-40, 1% SDS, 2 mM DTT, 5 mM NEM (N-ethylmaleimide), 3 mM iodoacetamide, protease and phosphatase inhibitors pH=7.3-7.4). For 5- 6 million cells 270 ul of ubiquitin IP buffer was added and incubated on ice for 10min. Cells were removed from the dish and transferred into tubes. 30 μl of 1X PBS was added and gently vortexed. Samples were boiled immediately for 20 minutes, until clear, then centrifuged at 14,000 rpm for 10 minutes. The boiled samples were diluted using IP Buffer without SDS to reduce the SDS concentration to 0.1%. For TRIM9 dimerization assay, HEK 293 cells transfected with Myc-tagged TRIM9 or TRIM9 variants and GFP tagged TRIM9 were lysed with RIPA buffer (50 mM Tris-HCl, ph 7.5, 150 mM NaCl, 1% NP40, 0.5% Sodium deoxycholate, 0.1% SDS with phosphatase and protease inhibitors).

For binding assays, all recombinant GST-tagged proteins were purified on sepharose-immobilized glutathione beads (ThermoScientific). For binding to endogenous TRIM67 or Ena/VASP, E15.5 mouse cortices were lysed in 0.5% NP40 lysis buffer (50 mM Tris pH7.5, 200 mM NaCl, 0.5% NP-40 with phosphatase and protease inhibitors). Lysates were pre-cleared with GST-glutathione-sepharose beads for 1 hour at 4°C with agitation and incubated with 5-10 μg of GST fusion protein or GST immobilized onto glutathione-sepharose beads at 4°C overnight. For binding of Myc-tagged TRIM67 variants, HEK293 cells were transfected and 24 hours later lysed in 1% NP40 lysis buffer and incubated overnight at 4°C with anti-myc antibody. For all binding assays, precipitated beads were washed three times with lysis buffer or PBS buffer and bound proteins were resolved by SDS-PAGE and analyzed by immunoblotting.

For co-immunoprecipitation assays, IgG-conjugated A/G beads (SCBT) were utilized to preclear lysates for 1.5 hours at 4°C with agitation. Myc antibody-conjugated A/G beads (SCBT) or Protein A/G beads (SCBT) coupled with a mouse anti-GFP Ab (Neuromab) or rabbit anti-VASP Ab (SCBT) were agitated within precleared lysates overnight at 4°C to precipitate target proteins. Beads were washed three times with lysis buffer and bound proteins were resolved by SDS-PAGE and analyzed by immunoblotting.

### Microscope Descriptions

All live cell images and immunofluorescence images were collected on an Olympus IX81-ZDC2 inverted microscope equipped with the following objective lenses: a UPLFLN 40x/1.39NA DIC objective (Olympus), UAPON 100x/1.49NA DIC TIRF objective (Olympus), a 20x/0.85NA UPlanSApo DIC objective lens (Olympus), and a 4x/0.13NA Plan Apochromat objective (Nikon), an automated XYZ stage (Prior) and an Andor iXon EM-CCD. Images were procured using the Metamorph acquisition software. Neuroanatomical images were acquired on an inverted laser scanning confocal microscope (Zeiss LSM 780, Zeiss) equipped with a 10X/0.4NA Plan Apochromat objective lens and 488-nm and 561-nm argon lasers. SIM images were obtained using a DeltaVision OMX SR imaging system (GE Healthcare) with a 60x 1.42 NA oil immersion objective in 2D-SIM super-resolution mode. SEM images were acquired using a Zeiss Supra 25 FESEM using a backscatter detector.

### Filopodia dynamics measurements

For filopodial dynamics measurements, wide-field epifluorescence images of mCherry were acquired every 2.5 seconds for 5-10 minutes. 600 ng/mL netrin-1 was added 40 minutes prior to imaging netrin-1 treated neurons. Filopodial protrusion and retraction rates and phase durations were measured from kymographs as the slope and duration of individual events, respectively. Filopodia lifetime was measured as the time from initial filopodial protrusion until retraction into the lamellipodial veil. Filopodial buckling and folding events were both counted manually; buckling events occurred when a filopodium collapsed following a breakage along the length of the filopodium. Folding events occurred when a filopodium collapsed into the veil sideways along its length as opposed to retracting perpendicularly.

### FRAP imaging

For FRAP assays neurons expressing GFP-VASP were imaged after 48 hours using 491 nm laser in TIRF mode every 0.5 seconds for 15 seconds, followed by a 1-second exposure with a solid state 405 nm laser in FRAP mode (bleach spot ~1.25 μm in diameter), followed by imaging every 0.5 seconds with the 491 laser in TIRF mode for 60 seconds. Netrin-1 treated filopodia were imaged within 5 minutes of addition of 400 ng/mL netrin-1. DUB-inhibited filopodia were imaged between 20 and 30 minutes of addition of 9 μM PR-619.

### Neuroanatomical imaging

All mice used for neuroanatomical studies were anesthetized on ice prior to decapitation, and heads were drop-fixed in 4% paraformaldehyde (PFA) for 72 hours. Heads were rinsed with 1x PBS twice for 24 hours prior to removal of and vibratome sectioning of the brains. For projection analysis in Nex-Cre/Tau^loxP-stop-loxP^GFP mice, 80 μm coronal sections were cut and every section was permeabilized in detergent solution (1x PBS + 0.1% Tx-100 + 0.2% Tween-20) for 1 hour on a shaker at RT. Sections were blocked in 10% BSA in 1x PBS for 5 hours, then placed in primary antibody solution (anti-GFP chicken (Aves ab1020 1:2000, in 1% BSA in PBS) for 24 hours on a shaker at 4°C. Primary antibodies were removed and sections were rinsed in 1x PBS for 1 hour prior to the addition of secondary antibody solution (AlexaFluor 488 chicken + 1% BSA in 1x PBS) for 24 hours on a covered shaker at 4°C. After post-secondary rinsing with 1x PBS, sections were mounted in TRIS/glycerol/n-propyl-gallate and were imaged with the 10x objective on the LSCM described above

### Axon Turning Assay

Micropass gradient devices were used to measure axon turning. Device preparation and experimental protocol is as described (Taylor et al., 2015). Sylgard 184 (Sigma) polymer was prepared as per manufacturer protocol and added to etched silicon wafers. Microfluidic devices were cut to the size of coverslips and fluid chambers were cut out, and devices were cleaned with tape, sprayed with 70% ethanol and allowed to dry for 45 minutes. Coverslips were applied to the face of devices with microfluidic grooves, and cell culture media was added to all chambers. After at least one hour, media was aspirated and *Trim67^+/+^* or *Trim67^-/-^* E15.5 cortical neurons were plated in devices. After axons entered the axon viewing area (2-4 days), a control gradient of dextran (starting at 1 μM) or gradient of netrin+dextran (600 ng/ml was established by adding solutions to one side chamber and removing media from the upper “sink” chamber. DIC (axons) and epifluorescence (dextran) images were acquired every 5 min for 8-18 hrs at 20x magnification.

## QUANTIFICATION AND STATISTICAL ANALYSIS

### Axon turning angle measurement

The angle of axon turning was determined as the angle between a line bisecting the growth cone after 18 hours of growth relative to the initial trajectory of the axon before the gradient, with positive angles indicating turning toward the source of the gradient. (Taylor et al., 2015).

### Colocalization and growth cone analysis

We quantified filopodial enrichment of endogenous TRIM67 distal 0.5 μm of the filopodia relative to the penultimate 0.5 μm of filopodia. We measured the proximity of TRIM67 to the filopodial tip by measuring the fluorescence ratio between the center and edge of a hypothetical Airy disc of a tip-localized punctum (tip localization ratio). This ratio should decrease as the distance of a fluorescent protein to the filopodial tip increases. Pearson’s correlation of colocalization between TRIM67 and Ena/VASP proteins was performed using regions of interest (ROI) drawn in filopodia and a Colocalization Test plugin for ImageJ with Fay randomization using images acquired with the 100x objective described above. Growth cone area, filopodia number, filopodia density and filopodia length were measured in ImageJ. Filopodia lengths were measured from the edge of lamellipodial veils to filopodia tips. For filopodia rescue assays, images of neurons expressing comparable levels of Myc(TRIM67 variants) were acquired with the 100x objective, and the number of growth cone filopodia recorded. TIRF imaging of TRIM67 and Ena/VASP proteins was performed after 48 hours *in vitro*, with the 100x objective and a solid state 491, 561 nm laser illumination at 100 nm penetration depth. Images were acquired every 0.5 seconds for 5 minutes.

### FRAP fluorescence recovery quantification

For analyzing FRAP imaging data, photobleaching was corrected by calculating an exponential decay from the last 30 seconds of imaging in a control region distant from the bleach spot (F = F_0_*e^-kt^, where F is fluorescence, F_0_ is initial fluorescence, k is the decay time constant, and t is time). Fluorescence recovery t_1/2_ and % were calculated from an inverse exponential decay (F = A*(1-e^-tt^), where F is fluorescence, A is recovery plateau fluorescence, t is the recovery time constant, and t is time). The % recovery was calculated as the plateau fluorescence divided by the average pre-bleach fluorescence, and t_1/2_ is the inverse of the recovery time constant τ. Both photobleaching and FRAP curves were fit to data using the Solver add-in of Microsoft Excel 2013.

### Western blot analysis

Western blots were quantified using ImageJ, with the total fluorescence of labeled bands representing relative protein which was then normalized to the control condition of each experiment. In the case of multiple controls in the same experiment (eg. VASP protein level quantification), results are normalized to the average of the controls on the same blot.

### Statistics

At least 3 independent experiments were performed for each assay. Data distribution normality was determined using the Shapiro-Wilk test. Normally distributed data were compared by unpaired t-test for two independent samples, or ANOVA with Tukey post-hoc correction, for >2 comparisons. For nonnormal data, the Mann-Whitney test was used or Kruskal-Wallis nonparametric ANOVA with Benjamini-Hochberg correction for >2 comparisons. Fisher’s exact test was used to compare growth cone morphology distributions. All data are presented as means +/-standard error of the mean (SEM) or as box plots (min, Q1, Q2, Q3, max) accompanied by all data points. Statistical significance was determined by an α of 0.05, and is represented as such: n.s. − p > 0.05, * p < 0.05, ** p < 0.01, *** p < 0.005.

## SUPPLEMENTAL INFORMATION

**Fig. S1:**
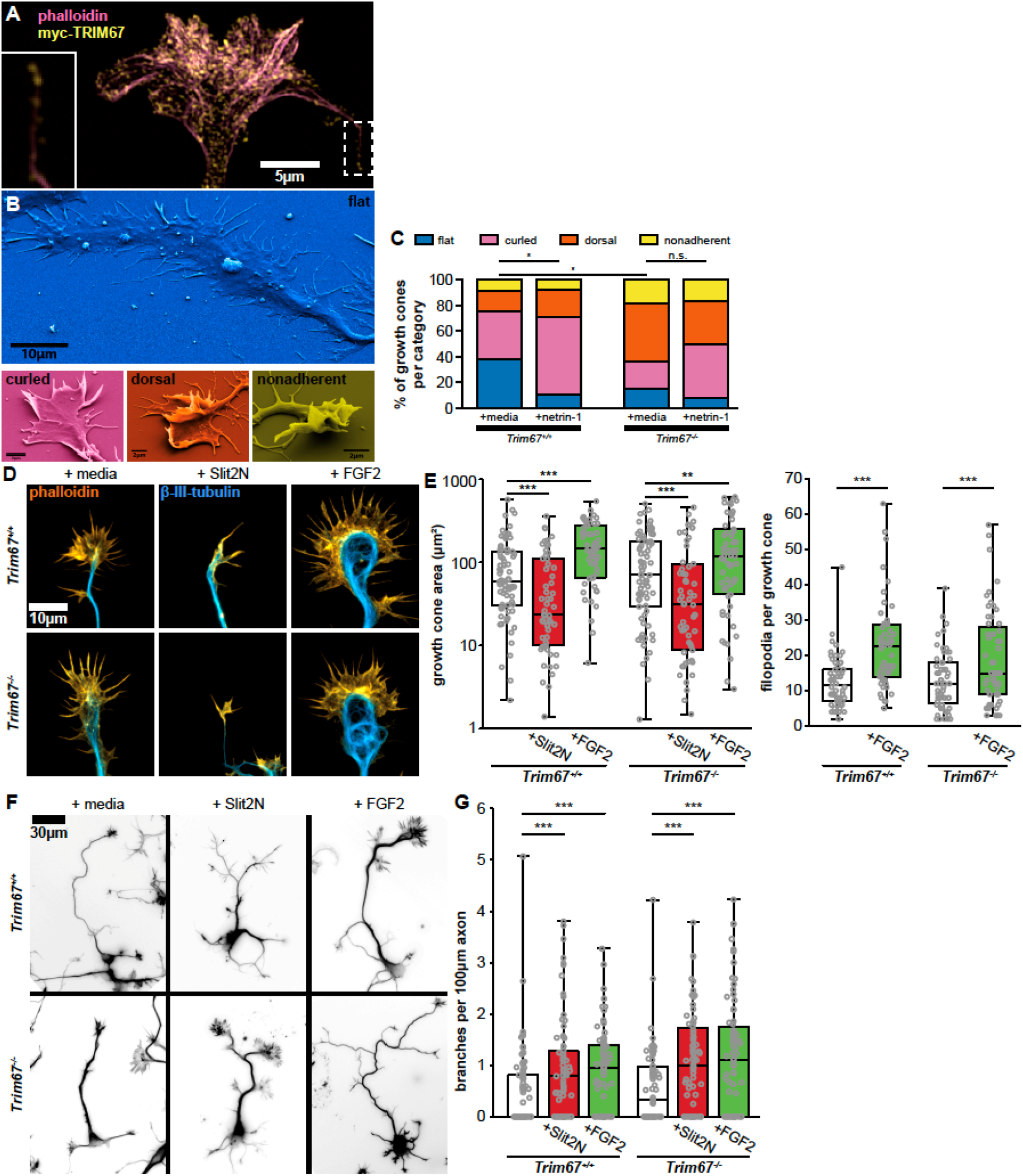
TRIM67 growth cone response to morphogens. **A)** Structured illumination microscopy image of myc-TRIM67 in an axonal growth cone, localizing to the tip of a filopodium (inset, same area as dashed box). **B)** Examples of growth cone morphological categories as identified by scanning electron microscopy. **C)** Quantification of growth cone morphology distributions of *Trim67^+/+^* and *Trim67^-/-^* cortical neurons treated with media or netrin-1. Distributions are compared by chi-square test. * - p < 0.05. **D)** Examples of cortical embryonic axonal growth cones treated with media, Slit2N, or FGF2. **E)** Quantification of growth cone area and filopodia per growth cone following 40 minutes of treatment with the indicated guidance cues. **F)** Inverted images of neurons combining staining of both filamentous actin (phalloidin) and β-III-tubulin, treated for 24 hours with media, Slit2N, or FGF2. **G)** Quantification of axon branching in response to the indicated guidance cues.

**Fig. S2:**
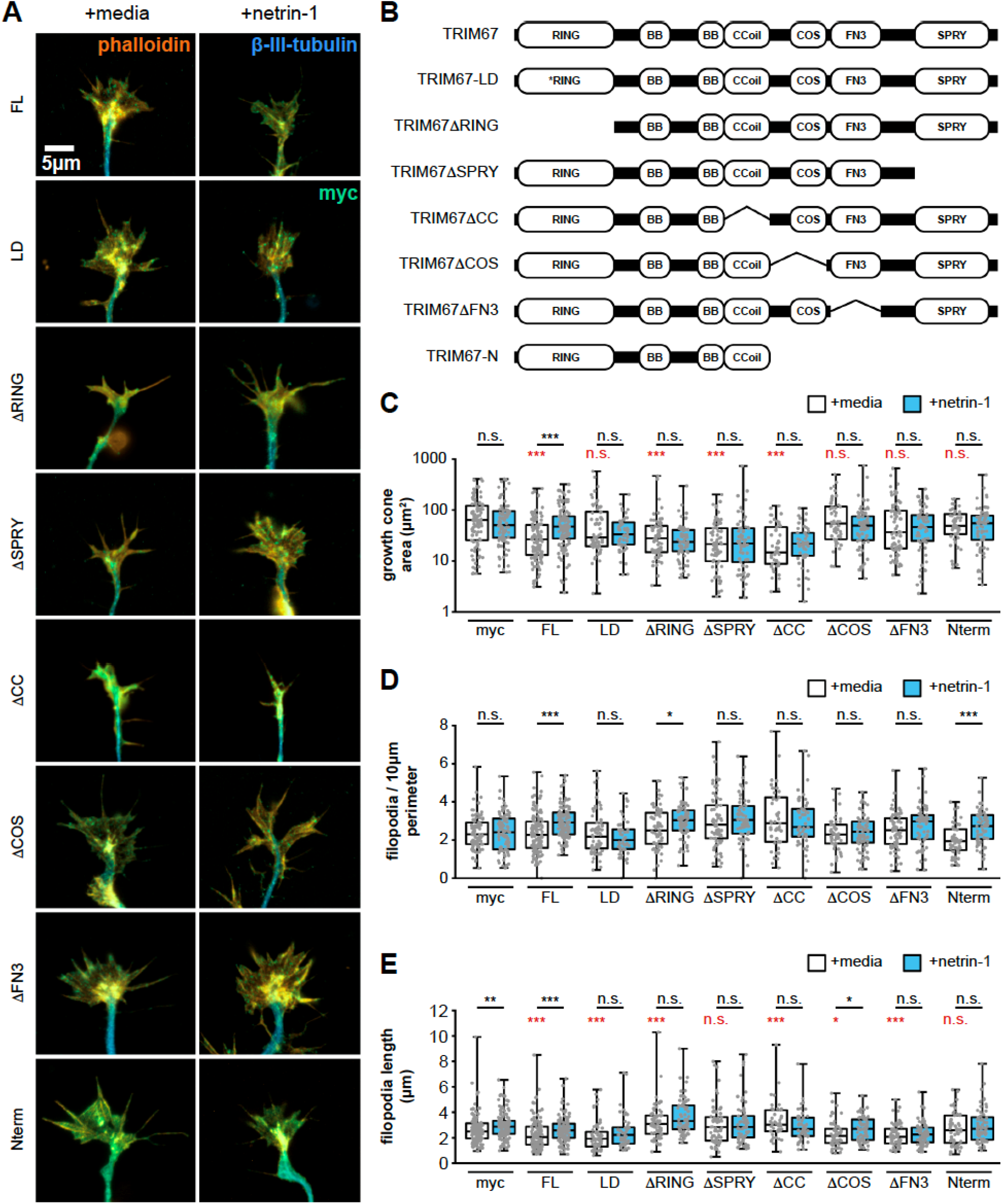
All domains of TRIM67 are required to fully rescue growth cone response to netrin-1. **A)** TRIM67 constructs used in structure-function assays. The RING domain of TRIM proteins contain zinc binding pockets necessary for E3 ubiquitin ligation activity and can mediate oligomerization of TRIM proteins (Freemont, 1993; Koliopoulos et al., 2016; Meroni and Diez-Roux, 2005); we therefore made both a RING-deletion construct (TRIM67ΔRING) and one containing mutations at cysteines 7 and 10 to abolish zinc binding in the RING domain and thus any ligase activity (TRIM67-LD). The coiled-coil (CC) domains of TRIM proteins mediate homo- and heterodimerization with other members of the same TRIM class (Sanchez et al., 2014; Short et al., 2002). In our previous investigation of TRIM9, the CC domain also interacted with the filopodial tip-localized actin polymerase VASP (Menon et al., 2015); therefore, we generated a construct of TRIM67 lacking the coiled-coil domain (TRIM67ΔCC). The C-terminal B30.2/SPRY domain of other TRIM proteins mediates interactions with binding partners (D’Cruz et al., 2013; Reymond et al., 2001), and in the case of TRIM9, interacts directly with the netrin-1 receptor DCC (Winkle et al., 2014), thus we made a construct of TRIM67 lacking this domain (TRIM67ΔSPRy). Binding to the microtubule cytoskeleton has been shown between the COS domain of the class 1 TRIM MID1 (TRIM18), and colocalization with tubulin has been shown with other class 1 members (Cainarca et al., 1999; Cox, 2012; Wright et al., 2016). The binding motif in the COS domain of MID1 is conserved throughout class 1 TRIMs (Wright et al., 2016), therefore we made a construct of TRIM67 lacking the COS domain (TRIM67ΔCOS). It has been suggested that the adjacent fibronectin type III (FN3) domain facilitates the interaction with microtubules, as a mutation in the FN3 domain of MID1 reduces association with microtubules (Cox, 2012; Perry et al., 1999; Schweiger et al., 1999); as such we made a construct of TRIM67 in which the FN3 domain was absent (TRIM67ΔFN3). Finally, we generated a construct possessing only the three N-terminal tripartite motif domains (TRIM67-N). All constructs possessed an N-terminal myc tag. **B)** ICC of *Trim67^-/-^* growth cones stained for actin, β-III-tubulin, and the indicated myc-TRIM67 construct, showing expression and distribution of each construct. **C)** Quantification of growth cone area, **D)** filopodial density, and E) filopodia length in neurons expressing each rescue construct. All statistical comparisons in red are to myc-expressing, untreated growth cones. * - p < 0.05, ** - p < 0.01, *** - p < 0.005.

**Fig. S3:**
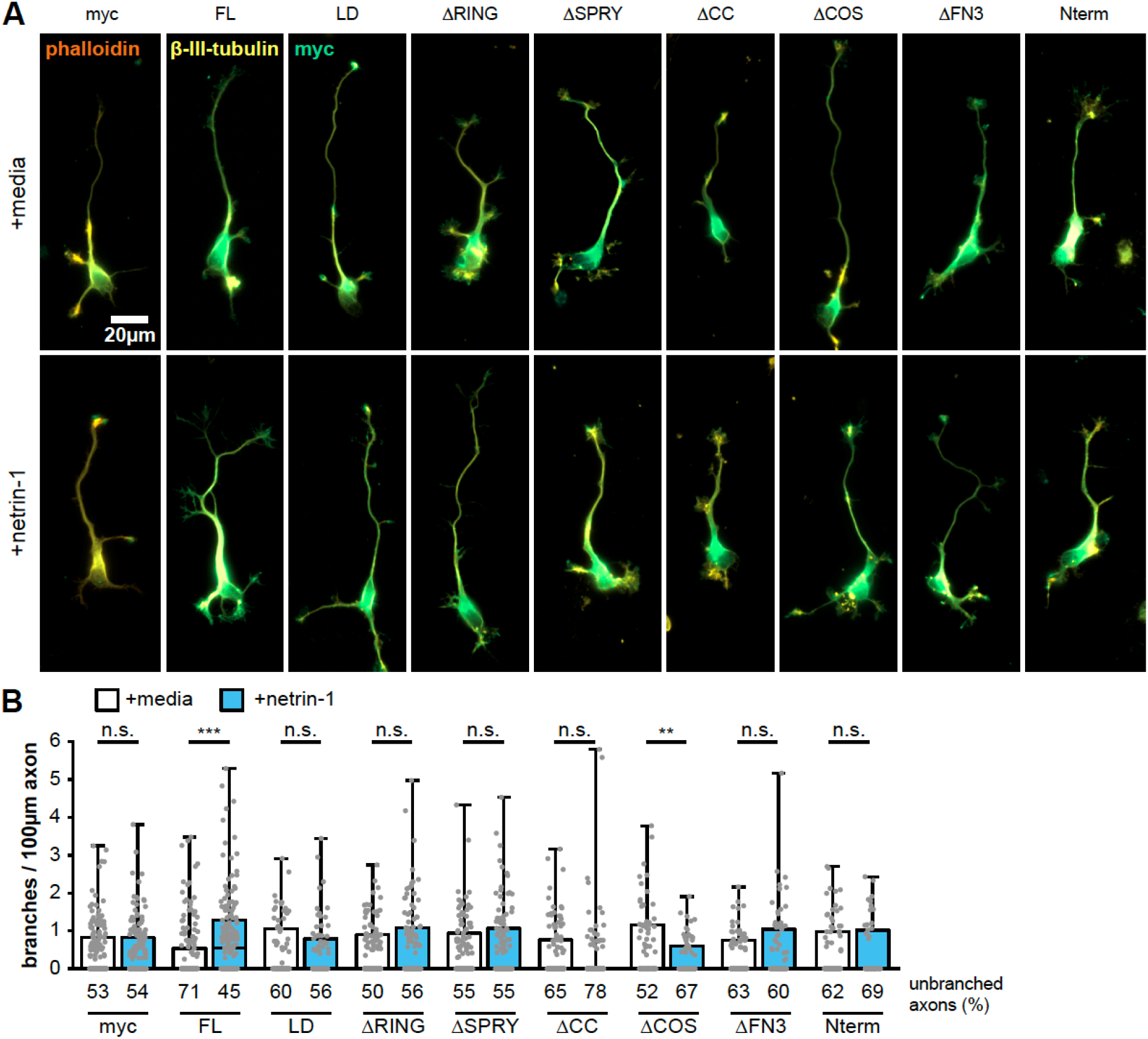
All domains of TRIM67 are required to rescue netrin-dependent axon branching. **A)** ICC of *Trim67^-/-^* neurons expressing indicated myc-tagged TRIM67 rescue construct, stained for myc (green), filamentous actin (phalloiding, red)and β-III-tubulin (yellow). **B)** Quantification of axon branches per 100 μm axon length. Percent of neurons with unbranched axons is shown below the x-axis. All statistical comparisons in red are to myc-expressing, untreated neurons. * - p < 0.05, ** - p < 0.01, *** - p < 0.005.

**Fig. S4:**
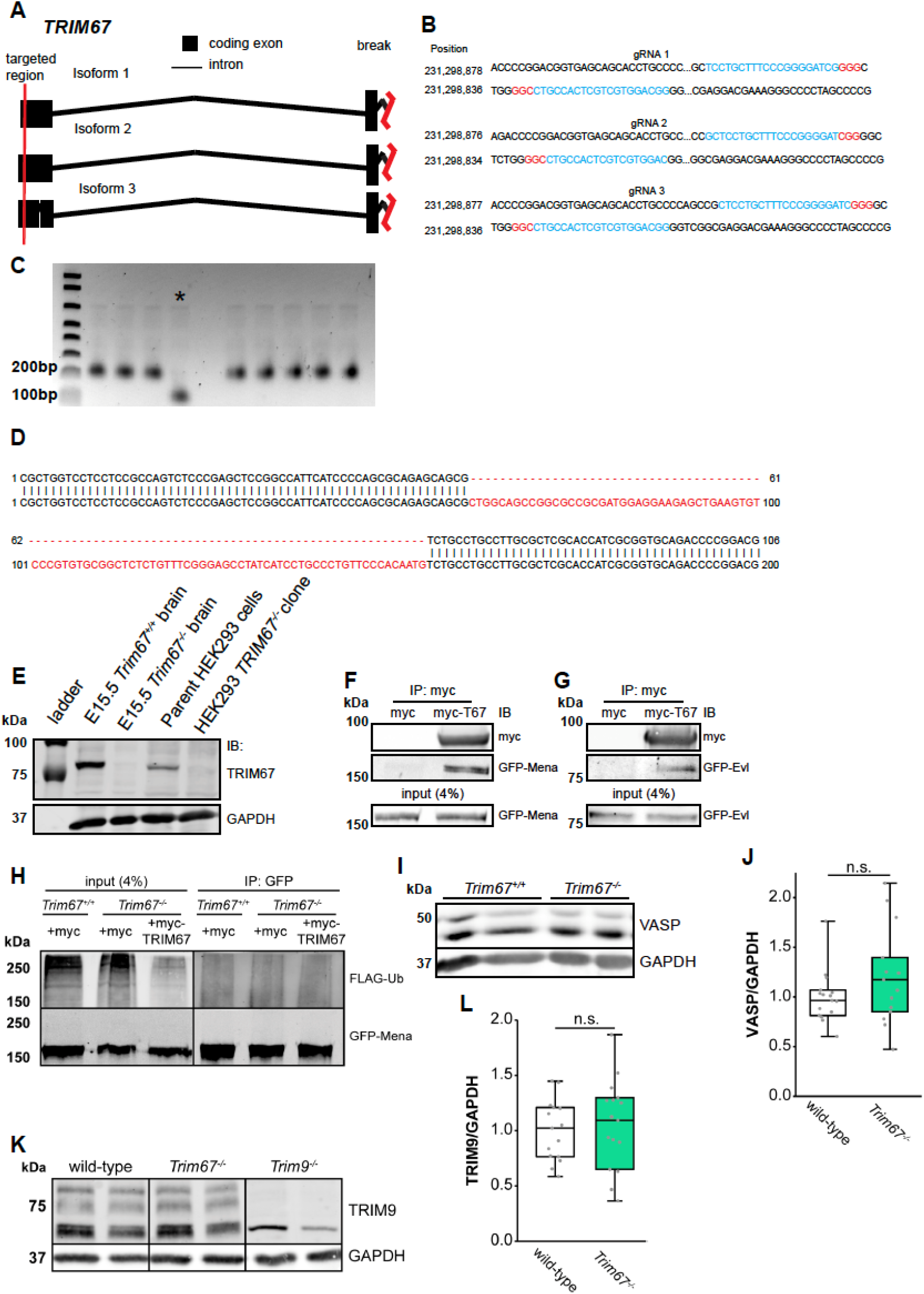
TRIM67 interacts with all members of the Ena/VASP family, and does not regulate VASP or TRIM9 protein levels. **A-E)** Generation of *TRIM67^-/-^* HEK293 cells lines. **A)** CRISPR gRNA were designed to target the first exon in all known isoforms of TRIM67. Red vertical lines indicate the regions targeted by the gRNAs. **B)** A set of three designed gRNAs (blue) targeting Exon 1 of TRIM67. Two complementary gRNAs per set were used with nickase-Cas9 to reduce off target effect potential. The NGG sequence is highlighted in red. **C)** PCR of genomic DNA of CRISPR clones indicates a deletion of ~ 100bp in CRISPR clone 3. **D)** Sequencing of HEK293 CRISPR clone 3 (top strand) shows a deletion of 94 base pairs in exon 1 of *TRIM67*. **E)** Immunoblot for TRIM67 and GAPDH in E15.5 murine brain lysate, *Trim67^-/-^* murine brain lysate, HEK293 lysates, and CRISPR-Cas9 clone 3 *TRIM67^-/-^* HEK293 lysates. **F)** Coimmunoprecipitation assay from *TRIM67^-/-^* HEK293T cells transfected with GFP-Mena and myc or myc-TRIM67. **G)** Similar coimmunoprecipitation with GFP-EVL. **H)** Immunoprecipitation of GFP-tagged Mena from denatured *TRIM67^-/-^* HEK293T cell lysate coexpressing FLAG-Ub and indicated myc or myc-TRIM67. **I)** Representative western blot of VASP in embryonic cortical lysate. **J)** VASP expression measured by Western blotting of embryonic cortical lysates, normalized to GAPDH. **K)** Representative western blot of TRIM9 in embryonic cortical lysate. Bottom band in *Trim9^-/-^* lysate is a non-specific band. L) TRIM9 expression from embryonic cortical lysates.

**Fig. S5:**
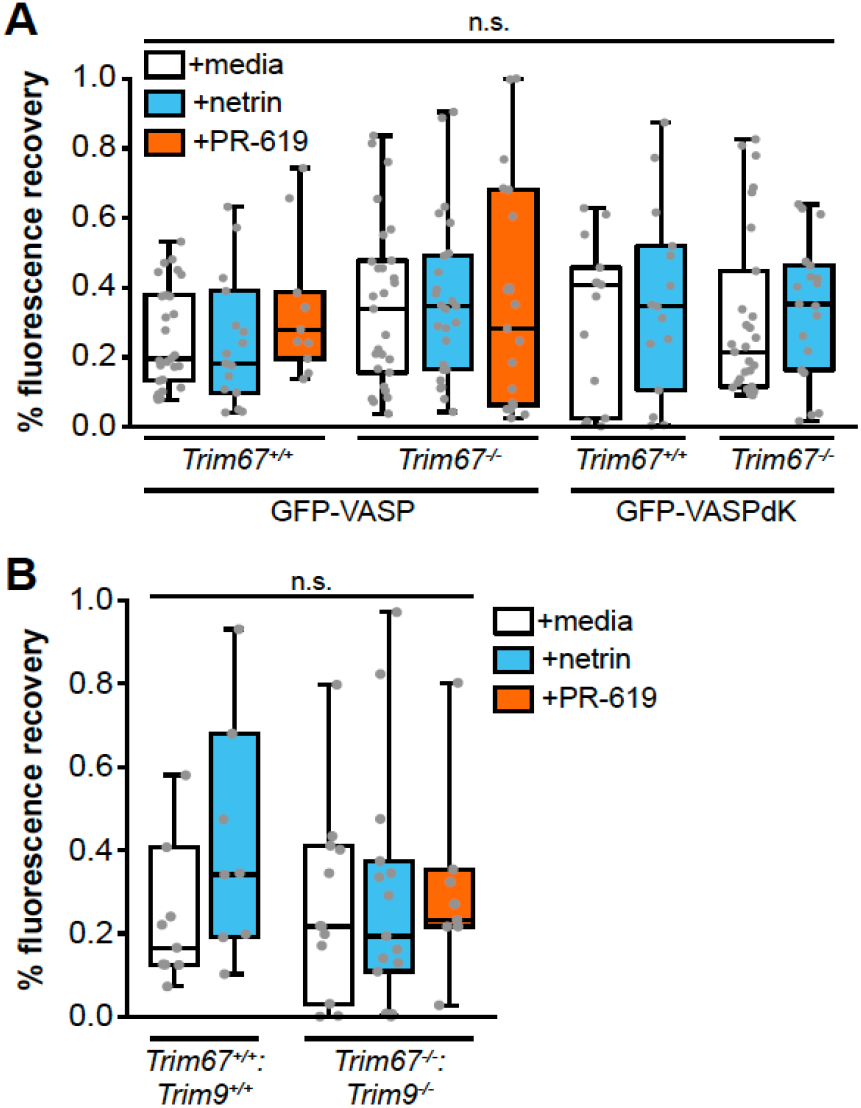
Percent recovery of fluorescence in FRAP assays. **A)** % recovery of fluorescence (mobile fraction) in FRAP assays reported in figure 5. **B)** % recovery of fluorescence in FRAP assays reported in figure 7.

**Movie S1: Live-cell imaging of filopodial dynamics of axonal growth cones**. Cultured embryonic cortical neurons expressing cytoplasmic mcherry, imaged every 2.5 seconds before or after 40 minutes of netrin-1 treatment. See Figure 4.

